# Spatial single-cell mass spectrometry defines zonation of the hepatocyte proteome

**DOI:** 10.1101/2022.12.03.518957

**Authors:** Florian A. Rosenberger, Marvin Thielert, Maximilian T. Strauss, Constantin Ammar, Sophia C. Mädler, Lisa Schweizer, Andreas Metousis, Patricia Skowronek, Maria Wahle, Janine Gote-Schniering, Anna Semenova, Herbert B. Schiller, Edwin Rodriguez, Thierry M. Nordmann, Andreas Mund, Matthias Mann

## Abstract

Single-cell proteomics by mass spectrometry (MS) is emerging as a powerful and unbiased method for the characterization of biological heterogeneity. So far, it has been limited to cultured cells, whereas an expansion of the method to complex tissues would greatly enhance biological insights. Here we describe single-cell Deep Visual Proteomics (scDVP), a technology that integrates high-content imaging, laser microdissection and multiplexed MS. scDVP resolves the context-dependent, spatial proteome of murine hepatocytes at a current depth of 1,700 proteins from a slice of a cell. Half of the proteome was differentially regulated in a spatial manner, with protein levels changing dramatically in proximity to the central vein. We applied machine learning to proteome classes and images, which subsequently inferred the spatial proteome from imaging data alone. scDVP is applicable to healthy and diseased tissues and complements other spatial proteomics or spatial omics technologies.

## INTRODUCTION

Mass spectrometry (MS)-based single-cell proteomics (scProteomics) has made tremendous progress within just a few years and can now quantify more than 1000 proteins in cultured cells ^1–3^. While this trajectory is promising, proteome depth, throughput and lack of spatial context limits biological use. We have recently introduced Deep Visual Proteomics (DVP), a spatial technology that combines imaging, cell segmentation, laser microdissection and MS into a single workflow to investigate complex tissues with various cell types and metabolic niches ^4^. DVP overcomes depth and throughput limitations with pooling the required number of cells with similar morphological features and staining patterns to identify statistically and analytically robust cellular phenotypes (“biological fractionation”). By its nature, it depends on prior knowledge of adequate markers of the cells of interest that resolve their heterogeneity. These markers might not be available for all subtypes of cells or those tissues that have rapidly changing proteome types such as heterogenous tumors. To address this, we here developed single-cell Deep Visual Proteomics (scDVP), a complementary technology that extends true scProteomics into the tissue context.

We use scDVP to explore spatial characteristics of hepatocyte subsets in mammalian liver - a highly organized and functionally repetitive tissue, in which the proteome of hepatocytes is determined by paracrine signaling, as well as oxygen and nutrient gradients ^5^. These metabolic gradients require distinct functional cell states along the portal to central vein axis. This phenomenon of liver zonation has been described by single-cell RNA sequencing for hepatocytes (scRNAseq) ^6,7^, FACS and MS-based proteomics ^8^, and multiplexed imaging ^9^. Despite this long and varied background, it still remains an open question how and to what degree hepatocyte proteomes differ spatially.

## RESULTS

### Robust isolation and characterization of hepatocyte shapes *in situ*

To map the proteome of mouse hepatocytes at single-cell resolution, we established a modular and automated workflow aimed at lossless sample preparation of the initial input cell for injection into the mass spectrometer (Fig. 1a). Mice livers were embedded and immediately frozen after cardiac arrest. We fixed 10 μm sections and stained them with a one-step protocol marking portal and central veins, the sinusoidal architecture, nuclei and cell membranes (Fig. 1b, see Methods). Individual cells were segmented by deep learning as before ^4^, and the resulting masks transferred to a laser microdissection microscope that automatically excised and collected individual shapes in 384-well plates. Given hepatocyte sizes of 20-30 μm, one shape cut from a 10 μm section corresponds to a third or half of a hepatocyte. We automated protein extraction and digestion by reagent addition into the same plate, omitting extra transfer steps, followed by peptide separation on the Evosep system and injection into a trapped ion mobility Time of Flight (timsTOF) SCP mass spectrometer ^10^ (Fig. 1a).

**Fig. 1:**
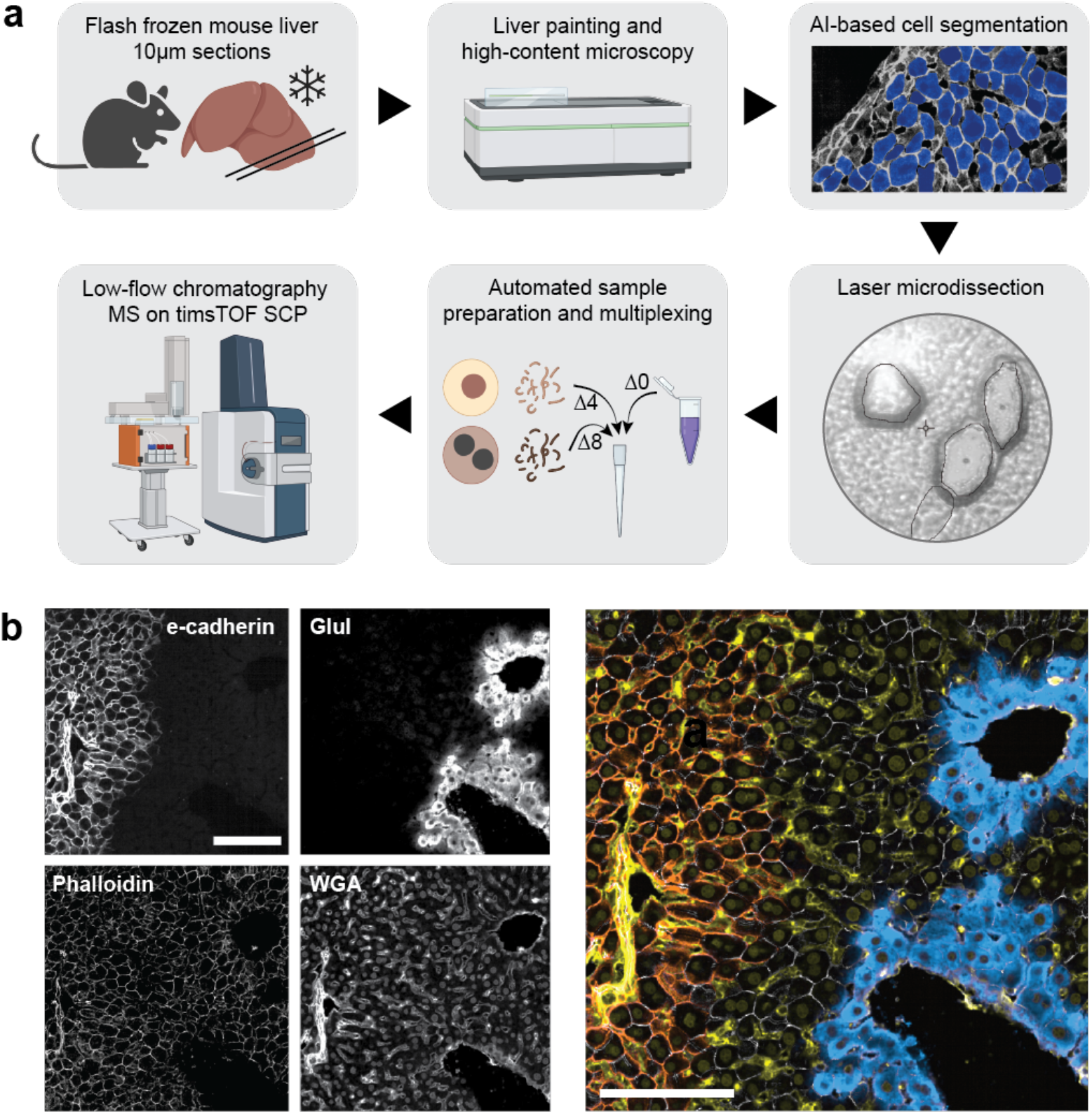
Isolation and characterization of individual hepatocyte shapes *in situ*. a, The scDVP workflow comprised embedding of fresh mouse liver tissue, staining and high-content microscopy, AI-guided hepatocyte segmentation, cutting and sorting of cells on a laser microdissection microscope, peptide preparation with or without dimethyl labeling. The Δ0 channel contains the reference proteome and Δ4 and Δ8 two individual samples, which are all analyzed by ultra-high sensitivity LC-MS. Created with BioRender.com. b, Liver painting with four stains. Left: E-cadherin marks portal vein regions, glutamate-ammonia ligase (Glul) surrounds the central vein, the cell segmentation marker Phalloidin, and the sinusoidal and nuclear counterstain wheat-germ agglutinin (WGA). Right: False color overlay of all channels. Scale bars 100 μm.

To establish an efficient workflow, we applied our established scProteomics protocol ^2^ and titrated the number of cells required to obtain a robust signal (Supplementary Fig. S1a). We performed initial experiments on five adjacent shapes per well (corresponding to about two complete hepatocyte cell masses), cut from randomly chosen locations. With these five shapes, we reached a median depth of 1,235 proteins across 230 samples (Supplementary Fig. S1b). Results confirmed expected liver biology, for instance by differential expression of the portal vein marker argininosuccinate lyase (Asl) and central Cyp2e1 (Supplementary Fig. S1c). Using zonation anchor proteins to arrange all samples in pseudo-space (Supplementary Fig. S1d), we characterized spatially enriched gene sets along the zonation axis. While oxidative phosphorylation and amino acid metabolism were among processes upregulated in proximity to the portal vein, drug metabolism and steroid hormone biosynthesis were increased proximal to the central vein, providing positive controls for low input proteomics (Supplementary Fig. S2a and S2b).

### Multiplex-DIA (mDIA) drastically increases proteome depth in single shapes

Encouraged by these spatial results, we next asked if single shapes alone could produce deep and interpretable proteomic results. To improve sensitivity further we adopted and optimized elements of our scProteomics workflow ^11^. These include addition of the surfactant n-Dodecyl-β-D-maltoside (DDM) to maximize peptide recovery ^12^, lowering the chromatographic flow rate to 100 nL/min for increased ionization efficiency ^2^ (‘Whisper gradients’ on the Evosep system) and achieving higher chromatographic resolution with zero dead volume columns (IonOpticks) ^13^. Most importantly, we added a labeled reference channel for multiplexed data-independent acquisition (DIA) that decouples identification and quantification ^11^ (Fig. 1a).

For scDVP, we constructed a dimethyl-labeled bulk liver reference. Our robotic sample preparation setup achieved greater than 99% labeling efficiency in all three channels (Supplementary Fig. S3a). We co-injected 10ng of the reference proteome together with the labeled proteomes of two single shapes. This resulted in a doubling of identified proteins with a median number of 1,726 proteins across three biological replicates and 455 single shapes, at twice the previous throughput (Fig. 2a). A maximum of more than 2,769 proteins were identified in one shape, and 3,738 unique proteins were found across all samples (Supplementary Fig. S3b, S3c). Four histone components ranked in the top 10 but we also found many transcription factors. The number of detected proteins correlated logarithmically with the microdissected area (Supplementary Fig. S3d), indicating that scDVP requires the highest possible MS sensitivity. Data completeness across all samples increased with median intensity per protein and coefficients of variation (CVs) were about 0.6 indicative of biological heterogeneity in the data (Supplementary Fig. S3e, S3f). We hypothesized that the nuclear proportion in the cell slice would correlate with the intensity of these histones. Indeed, shapes with lowest histone intensities did not have any evident nuclear signal, while top intensities were in shapes with large or two nuclei. In addition to this, the intensity of the top four abundant histone proteins was highest in arterioles that we cut as technical control structures that are composed of more than one cell and nucleus (Fig. 2b).

**Fig. 2:**
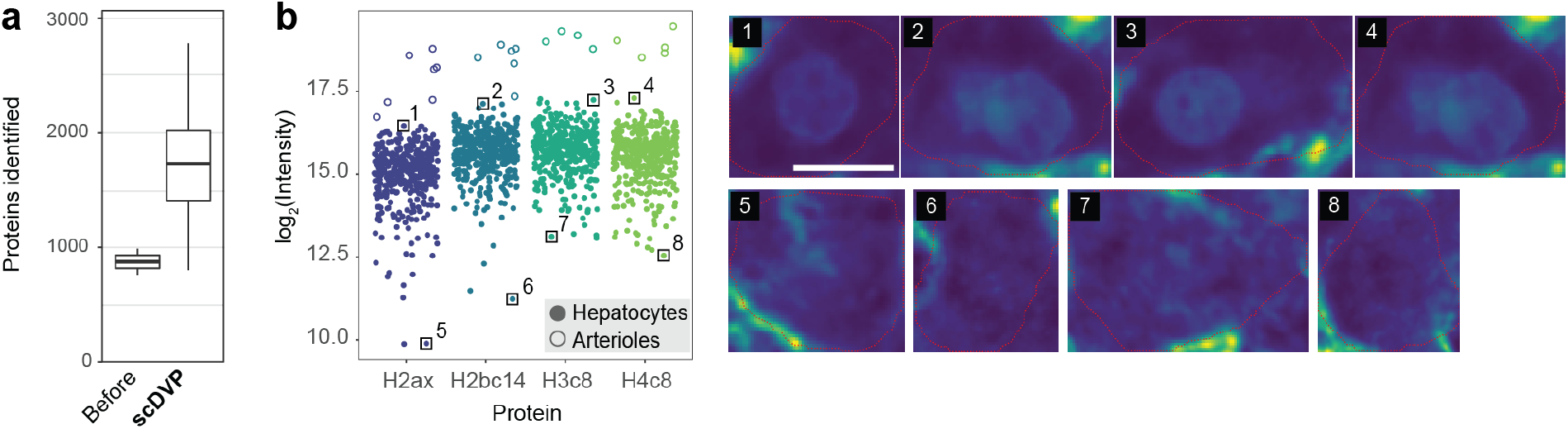
Depth of single shape proteomes and estimation of the nuclear compartment. a, Number of proteins quantified in the original workflow (single shape, 44 min Evosep gradient, 15 cm column at 500 nL/min, dia-PASEF ^14^ without optimized windows, library-dependent search in DIA-NN ^15^; numbers identical to Supplementary Fig. S1A), versus our new mDIA (two single shapes and reference proteome, 31 min Evosep gradient, 15 cm column at 100 nL/min, dia-PASEF with optimized window design, library-dependent search in DIA-NN). b, Left: Intensity of the top four histone proteins across all samples, including hepatocytes and quality control arteriole structures. Right: WGA-stain of cells corresponding to marked data points in the scatterplot. Scale bar: 10μm.

### Single shape proteomes accurately reflect hepatocyte zonation

To test the biological validity of our proteomics data, we first reduced dimensionality in a principal component analysis (PCA) which revealed that PC1 represented the measured distance of a hepatocyte to portal and central vein (Fig. 3a and 3b). Overlays of known liver zonation markers including Cyp2e1 and Asl showed opposite visual enrichment along PC1 (Supplementary Figs. S4a and S4b). In contrast, PC2 Eigenvalues did not correlate with measured distance or hepatocyte zonation markers but rather with cytoskeletal components (Supplementary Figs. S4c and S4d). PC2 was also the dimension in which portal arterioles, which we excised as technical controls, separated from hepatocytes (Supplementary Fig. S4e). Based on PC1 Eigenvalues we grouped the data into eight bins, which was a good compromise between meaningful separation and a sufficient number of samples per group. ANOVA testing revealed that 53% of all proteins detected in at least half of the samples were significantly different between zones (FDR < 0.05, Supplementary Fig. S5a). Zonation was also apparent after PC1 Eigenvalue sorting at the total proteome level (Fig. 3c) and for known hepatocyte zonation markers (Fig. 3d). Only 5.8% of these proteins were expressed equally in all zones (multiple testing adjusted Shapiro-Wilk test, p > 0.05), including Electron Transfer Flavoprotein β (Etfb), the electron acceptor in mitochondrial fatty acid β-oxidation (Supplementary Fig. S5b).

**Fig. 3.**
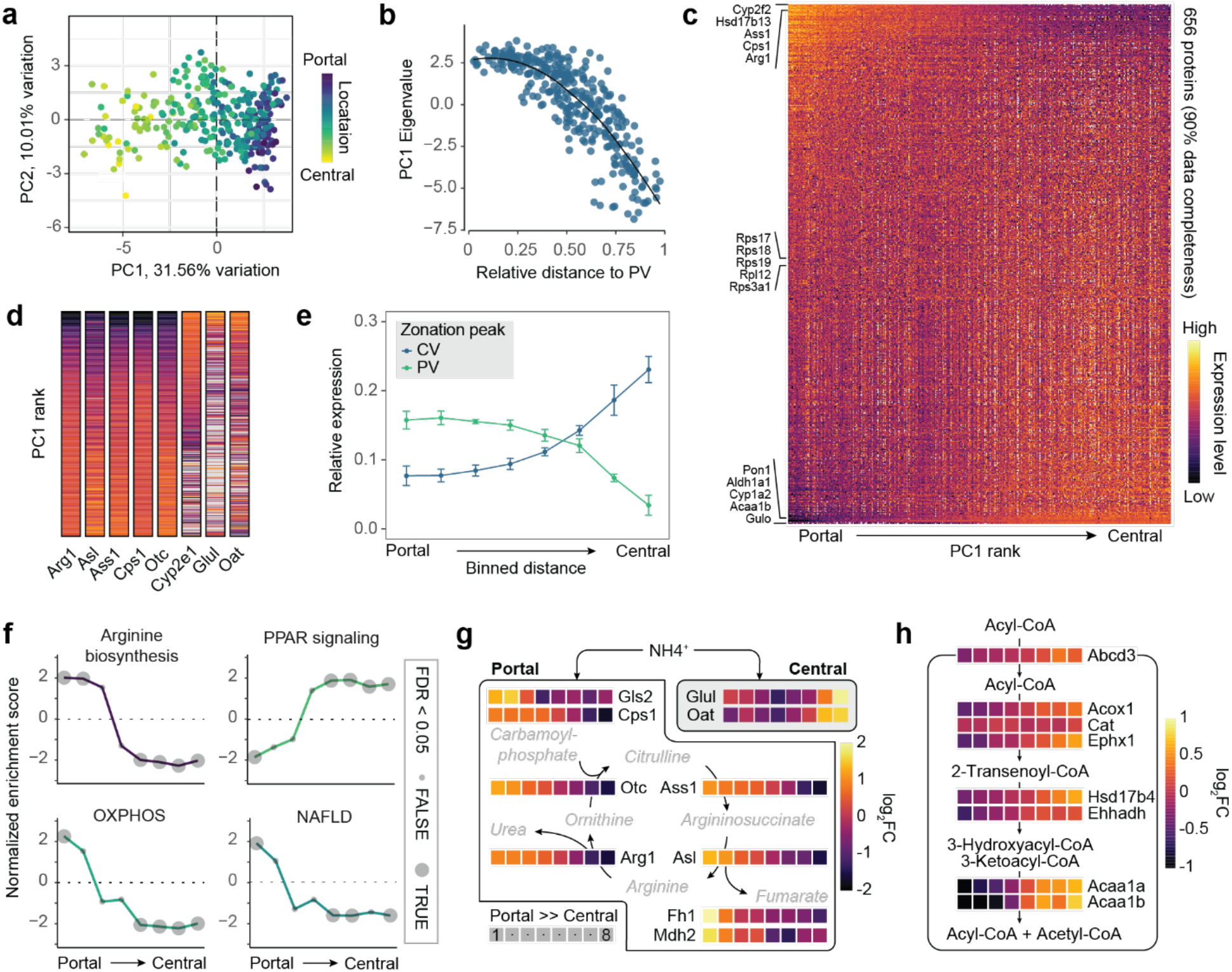
Single shape proteomes are accurate descriptors of zonated hepatocytes. **a,** Principal component analysis (PCA) of all hepatocytes. The color overlay corresponds to the ratio of measured distance portal vein over central vein in the microscopy image. **b,** Measured distance ratio versus Eigenvalues of PC 1. Relative distance of 0 is at the portal vein, of 1 is at the central vein. Black: Smoothing curve. **c,** Heatmap of protein expression as z-score per protein across all samples. Proteins are ordered according to ANOVA fold change across eight PC1-guided bins; samples are ranked according to their PC1 Eigenvalue. The five top and bottom proteins are given as well as five ribosomal subunits within the 10 middle ranks. Only proteins that were detected in 90% of all samples are included. **d,** Protein expression as in c of selected marker proteins. **e,** Expression of the top 10 significant proteins in eight measured distance bins, relative to total expression from portal to central. Zonation peak at PV: positive ANOVA fold change (n = 6), and vice versa (n = 4). Error bars represent standard deviation. **f,** Selected gene sets in individual PC1-guided bins versus all others bins, depicting normalized enrichment score after gene set enrichment analysis. Dot size: significance. PPAR: Peroxisome proliferator-activated receptor; OXPHOS: Oxidative phosphorylation; NAFLD: Non-alcoholic fatty liver disease. **g,** Levels of urea cycle and connected enzymes from portal (left) to central (right) PC1-guided bins. Portal box: active in portal regions. Central box: active in central region. **h,** Levels of peroxisomal enzymes related to very-long chain fatty acid degradation, spatially resolved as in g.

The correlation of zonal proteomes indicated that portal and periportal regions were more similar to one another than central and pericentral zones (Fig. 3b, Supplementary Fig. S5c). Indeed, the spatial expression of the top-10 significant zonation markers followed a hockey-stick curve from portal to central (Fig.3e), similar to Wnt-controlled transcripts in a scRNAseq dataset ^6^ and in line with a central vein origin of Wnt signaling ^16^. In contrast, this pattern was absent for the hits with the highest p values (least zonated hits, Supplementary Fig. S5d).

A cross-omics comparison with scRNAseq data ^6^ confirmed the directionality of the most prominent zonation markers (Pearson’s R 0.85, Supplementary Fig. S6a and S6b). However, the relative expression differences from portal to central vein were less extreme in our proteomics dataset compared to scRNAseq. Notably, a number of proteins were regulated only in the RNA or protein dimension, or even inversely correlated (Supplementary Fig. S6c and S6d), such as Epoxide Hydrolase 2 in the peroxisomal fatty acid degradation pathway. Members of glutathione metabolism had similar spatial distribution in both datasets (Supplementary Fig. S6e).

Enrichment of functional protein sets across PC1-guided bins confirmed that arginine biosynthesis and oxidative phosphorylation were highly enriched towards the portal vein (Fig. 3f). In line with this, all proteins participating in ammonia fixation of the urea cycle were portally expressed, while ammonia-capture on glutamate were strongly central (Fig. 3g). To our surprise, several other disease and signaling related pathways were also zonated including those involved in Non-Alcoholic Fatty Liver Disease (NAFLD) and Peroxisome proliferator-activated receptor (PPAR) signaling (Fig. 3f). This was corroborated by prominent central expression of enzymes required for peroxisomal degradation of very-long-chain fatty acids, and ω-oxidation of dicarboxylic C12 fatty acids such as coconut oil (Fig. 3h). We conclude that the spatial proteome data from single hepatocyte shapes is biologically accurate and informative.

### Proteomes of single hepatocytes are regulated by spatial context

Combining the single shape proteomes with their inherent spatial information and staining intensities, scDVP revealed clear dependence of fluorescent intensities with the eight proteome classes established above (Fig. 4a and 4b). We compared proteomes, in which cells were direct neighbors (n = 26) to pairs of not neighboring cells that were assigned to the same proteomic bin.

**Fig. 4.**
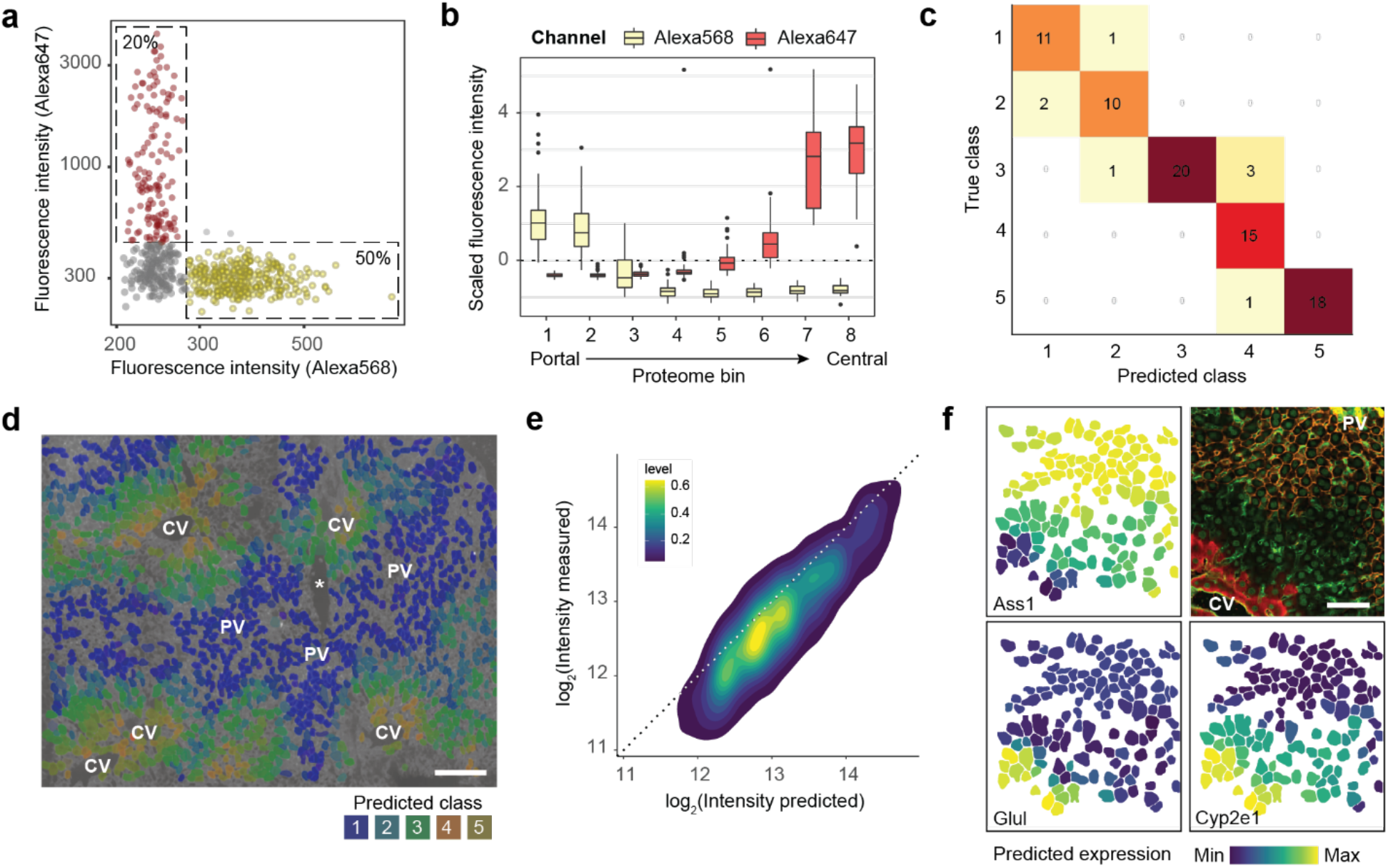
Combining imaging and proteome data for a machine-learned model. **a,** Fluorescence intensities of Alexa568 (portal vein marker E-cadherin) and Alexa647 (central vein marker Glul), with percentages in indicated bins. **b**, Intensities of the spatial markers as in (a) across eight PC1-guided proteome bins. **c**, Confusion matrix of a machine-learned model with five classes, informed by microscopy and proteomics data. **d**, Predicted classes of segmented hepatocytes. Hue is maximum class probability. **e**, Density plot of predicted versus measured intensities of a section excluded from machine learning (R = 0.78). **f**, Spatial depiction of data in (e) with microscopy ground truth on top right, and three predictions. CV: Central vein; PV: Portal vein; *sectioning artifact. Scale bar 50 μm.

Neighbors had significantly more similar intensity profiles, yet only for portal but not central markers (Supplementary Fig. S7a). Interestingly, periportal cells that were assigned to the same proteome class were typically multiples of 350 μm apart within one biological replicate (Supplementary Fig. S7b, S7c), which corresponds to the expected vein-to-vein distance in mouse liver tissue. This also held true for cells of the intermediate and central zones, although less prominent.

Encouraged by the evident complementarity between extensive proteomics and spatial data, we reasoned that the microscopic image could contain sufficient information to predict the proteome. To this end, we trained a machine learning model on 17 features to predict the proteome classes from imaging data. We grouped the training set into five proteome classes by k-means clustering (Supplementary Fig. S8a), and used the information in all imaging channels as predictors (Supplementary Fig. S8b). This model reached an average precision of 0.94 (Supplementary Fig. S8c and S8d), correctly assigning the proteome class of almost all cells. Errors occurred exclusively between spatially neighboring classes (Fig. 4c).

We tested model performance on a new section (not used in training) from which we measured 60 single-shape proteomes. Visual inspection indicated that the predicted classes were correctly located in proximity to central or portal vein, even in the presence of cutting artifacts (Fig. 4d). We used the class probabilities as weights to predict the spatial proteome, which accurately approximated overall protein intensities (R = 0.78 between prediction and measurement, Fig. 4e). When predicting the proteome of a larger section for all quantified proteins the ML model correctly assigned the spatial directionality of zonation markers, as well as their expected extension into the intermediate zone (Fig. 4f). Thus, the model confirms the accuracy of measured single shape proteomes, and is furthermore a potent predictor of spatial proteomes across any imaged areas.

## DISCUSSION

Here we present a single-cell, spatial map of the murine liver acquired by MS-based proteomics. Our approach successfully combined microscopic imaging data with ultra-high sensitivity proteomics, building on four major technological advances: (1) AI-assisted segmentation and laser microdissection, (2) multiplex-DIA, (3) low-flow gradients, and (4) the ultra-high sensitivity of a timsTOF SCP mass spectrometer.

To date MS-based single-cell proteomics has been exclusively reported for cell suspensions. State-of-the-art workflows currently reach a proteomic depth of up to 2,000 proteins in cultured cells, with about 250 pg of cellular protein mass. This is similar to the protein material in our sliced hepatocytes taking the section thickness of 10 μm and hepatocyte size of 20-30 μm into account. With our scDVP workflow, we achieved more than 1,700 proteins per single shape (and up to 2,700) despite working from sections that were fixed, stained, imaged and laser dissected. The size of our shapes correlated strongly with the number of identified proteins, suggesting that scDVP is currently limited by MS sensitivity and will thus profit from continuous technical developments.

Our proteomics data from single shapes correctly and accurately recapitulates liver physiology by direction, extent and spatial organization of zonation. More than half of quantified proteins were significantly different between portal and central zone, in line with scRNAseq data ^6,17^. The fact that we detected all of the previously used markers of liver zonation ^6^, suggests that our proteomic depth is sufficient to integrate into other omics datasets. This became further apparent on the level of functional pathways, including signaling and disease pathways. Interestingly, peroxisomal degradation of very-long-chain fatty acids, as well as dicarboxylic C12 fatty acids such as coconut oil, was enriched in proximity to the central vein. Biochemical evidence by radiolabeling experiments support the notion that non-mitochondrial fatty acid oxidation localizes to pericentral regions ^18^. A rhythmic expression pattern has been previously shown for a large number of liver transcripts and proteins ^17,19^. While we have not covered the temporal aspect here, the scDVP approach could contribute to such studies by adding a spatial dimension.

In the previously described DVP workflow we used pools of cells combined on the basis of common features, such as the expression intensity of already known markers, or morphology ^4^. This approach allows a deep, rapid and robust proteome characterization that accurately represents the underlying biology. By analyzing single cellular shapes without prior assumptions, scDVP now removes the dependency on established markers or features. This makes it a promising approach in heterogeneous tissues with partially or not defined subtypes of cells, such as in many tumor tissues. Moreover, scDVP can be a method of choice to map proteomic disturbances along gradients of, for instance, signaling factors, nutrients or gases, and in physiological settings that may create impediments for other omics methods, for instance in extracellular fibrotic scars.

We have shown that single-cell data can be used to train an accurate machine learning model that predicts the proteome class from visual information only. Evidence suggests that morphological features such as nuclear vacuolation and texture associate with zonation, and can even serve as a progression and stratification marker of non-alcoholic fatty liver disease ^20^. Combining such easily available features and extensive proteomic sampling can clearly lead to higher precision of the predictive models. Transfer learning might then extend the approach to many new areas, as already shown for single-cell transcriptomics data ^21^. The modular nature of scDVP makes it compatible with other spatial omics technologies such as spatial transcriptomics, epigenomics ^22^ or multiplexed imaging. In conclusion, ScDVP is a powerful tool for basic discovery science, working in concert with DVP and other omics methods to enrich spatial workflows.

## ACKNOWLEDGEMENTS

We thank our colleagues at the Department of Proteomics and Signal Transduction at the Max Planck Institute of Biochemistry as well as our colleagues at the Center for Proteome Research in Copenhagen for their input and support. We are particularly grateful for input and help from Katherine Madden, Isabelle Bludau, Andreas-David Brunner and Leonie Zeitler. FAS is an EMBO postdoctoral fellow (ALTF 399-2021). SCM is a PhD fellow of the Boehringer Ingelheim Fonds. JGS has received funding from the European Respiratory Society and the European Union’s H2020 research and innovation program under the Marie Sklodowska-Curie RESPIRE4 grant agreement No 847462. This study has been supported by the Horizon-2020 under the MICROB-PREDICT program (No 825694) and ISLET (No. 874839), by the Max-Planck Society for Advancement of Science, by the Chan Zuckerberg Initiative (CZF2019-002448), and by grants from the Novo Nordisk Foundation, Denmark (grant agreement NNF14CC0001 and NNF15CC0001).

## CONTRIBUTIONS

FAR, MTh and MM conceptualized the scDVP workflow. MTh, FAR, PS and MW acquired mass spectrometry data. FAR, MTS, LS, and AMe performed data analysis. MTS trained the machine-learned model. CA developed quantification software. SCM optimized the high-content microscopy pipeline. FAR, MTh, LS, AMe, SCM, ER, TMN, and AMu developed and optimized the experimental scDVP workflow. JGS, AS and HBS provided mouse samples. FAR, MTS, PS and TMN curated data. MM and FAR supervised the project. FAR and MM wrote the original manuscript draft. All authors read, revised and approved the manuscript.

## METHODS

### Mouse experiments and organ harvesting

Pathogen-free male and female 10-week-old C57BL/6J-rj mice were purchased from Janvier (France) and maintained at the appropriate biosafety level under constant temperature and humidity conditions with a 12 h light cycle. Animals were allowed food and water ad libitum. All experiments were performed on 12-or 13-week-old wild-type mice. These were sacrificed by cervical dislocation, and the liver was rapidly excised through a ventral opening of the peritoneum. The organ was rinsed in cold PBS, and the left lateral lobe was divided into three pieces. For this study, the distal-caudal quarter was embedded in Optimal Cutting Temperature (O.C.T.) medium (Sakura Finetek, Japan) in 15 mm disposable cryomolds (Sakura Finetek, Japan) and frozen in isopentane that was prior cooled to dew point in liquid nitrogen. Fully solidified blocks were transferred to dry ice, and then to a -80 °C freezer until further processing. Animal handling and organ withdrawal were performed in accordance with the governmental and international animal welfare guidelines and ethical oversight by the local government for the administrative region of Upper Bavaria (Germany), registered under ROB-55.2-2532.Vet_02-16-208.

### Immunofluorescence staining

Two-micrometer polyethylene naphthalate (PEN) membrane slides were pre-treated by UV ionization for one hour at 254 nm. Without delay, slides were consecutively washed for five minutes each in 350 mL acetone, 7 mL VECTABOND reagent to 350 mL with acetone, and then washed in ddH2O for 30 seconds before drying in a gentle nitrogen air flow. For sectioning, tissue blocks were transferred to a cryostat (Leica CM3050) at -18 °C chamber and -15 °C object temperature, and left to equilibrate for 30 minutes. Blocks were then trimmed, and final sections were cut at 10 μm thickness with a disposable high-profile blade (Leica 818). Frozen sections were transferred to pre-treated, cold PEN-membrane slides, and melted for less than 5 seconds on a room temperature surface. The sections were then fixed in pre-warmed 4% PFA in PBS at 37 °C, then in 95% ethanol at room temperature, and finally again in 4% PFA in PBS at 37 °C. Slides were rinsed in PBS and left in 5% BSA-PBS blocking solution for one hour until staining. Sections were stained for one hour at 37 °C in a humid and dark chamber with 200 μL of a one-step liver painting in 1% BSA: 1:300 phalloidin coupled to Atto-425 (Sigma 66939), 1:200 wheat-germ agglutinin (WGA) coupled to Alexa Fluor 488 (Invitrogen W11261), 1:100 anti-e cadherin coupled to Alexa Fluor 555 (BD 560064), anti-glutamine synthase (Abcam ab176562), and 1:500 anti-rabbit nanobody coupled to Alexa Fluor 647 (Chromotek srbAF647-1-100). Slides were washed three times for two minutes in PBS in the dark, and mounted with 21 μL ProLong Diamond mounting medium (Invitrogen, P36961) and a 22 × 22 mm #1.5 coverslip. Slides were stored until imaging in 50 mL tubes with desiccating material at 4 °C.

### High-content imaging

Sections were imaged on an OperaPhenix high-content microscope, controlled with Harmony v4.9 software, at 40X magnification, binning of two and a per tile overlap of 10%. At an excitation wavelength of 425 nm, 555 nm and 647 nm, 80% laser intensity were used at an illumination time of 100 ms, while 20% and 20 ms were used in the 488 nm channel. E-cadherin and glutamine synthetase were imaged simultaneously, while phalloidin and WGA were imaged consecutively.

### Image post-processing

Acquired images were flat-field corrected using the Harmony software. Stitching of image tiles was performed using the ashlar python API ^23^ with a max shift value of 30. Stitched images were exported as .tif files and imported into the Biological Image Analysis Software (BIAS) ^4^ with the packaged import tool. In BIAS, large tif images were first retiled to 1024 × 1024 px at an overlap of 5%. Hepatocytes were identified with a deep neural network for histological cytoplasm segmentation on the basis of CFP staining at 1.2 input spatial scaling, 40% detection confidence and 30% contour confidence. Only contours between 30 μm^2^ and 300 μm^2^ were taken into consideration. After removal of duplicates and false identifications by supervised machine learning, contours were exported together with three calibration points that were chosen at characteristic tissue positions. Contour outlines were simplified by removing 99% of data points. For five-shape proteomes, directly adjacent shapes forming a pentagon-like structure were manually picked. Single shapes were randomly picked and every 15^th^ to 25^th^ shape was assigned to adjacent wells in a 384-well plate. Arterioles were manually assigned based on WGA signal, ellipticity, and proximity to the E-cadherin positive portal vein.

### Laser microdissection

Contour outlines were imported after reference point alignment, and shapes were cut by laser microdissection with the LMD7 (Leica) in a semi-automated manner at the following settings: power 59, aperture 1, speed 60, middle pulse count 1, final pulse -1, head current 48 - 52%, pulse frequency 3282, offset 100. For the five-shape experiment, the microscope was controlled with LMD v8.2, with which five directly adjacent shapes were sorted into a low-binding 384-well plate (Eppendorf 0030129547) with one empty well between samples. Single shapes were cut and sorted with the software LMD beta 10 after calibration of the gravitational stage shift into 384-well plates into all wells, leaving the outermost rows and columns empty. Plates were sealed, centrifuged at 1,000xg for 5 minutes and then frozen at -20 °C until further processing.

### Reference peptide preparation for five-shape and single-shape proteomes

The proximal part of two biologically independent lobes of the same mice as in the scDVP experiments was used to construct a library. The tissue embedded in O.C.T. was removed from -80 °C and directly disintegrated in a plastic bag with a rubber hammer. Pieces of approximately 1mm^3^ were transferred into a low-binding 96-well plate with magnets (BeatBox Tissue Kit, Preomics, Germany), covered with 50 μL of 60 mM triethylammonium bicarbonate buffer with 10% acetonitrile (ACN; lysis buffer), and lysed in a BeatBox (Preomics, Germany) at standard settings for 10 minutes. Samples were then boiled at 96 °C for 20 minutes, transferred to 1.5 mL low-binding tubes, filled up to 500 μL with lysis buffer and sonicated for 5 times 30 seconds on/off cycles. After centrifugation at 2,000xg for 1 minute, the protein concentration in the supernatant was estimated on a Nanodrop, and LysC and trypsin were added at a protein-to-enzyme ratio of 1:100. After digest for 20 hours, samples were acidified to 1% TFA, centrifuged at 3,000xg for 10 minutes at room temperature, and dried in a SpeedVac for 30 minutes. Digest was filled to 1mL with buffer A (0.1% formic acid [FA]), and desalted on C-18 columns (Waters WAT036820). They were activated and equilibrated with 2mL of methanol, 2 mL of buffer B (100% ACN, 0.1% FA) and 2mL of buffer A, before sample loading. Peptides were washed with buffer A two times, eluted in 80% ACN with 0.2% FA, and dried down.

### Library fractionation for five-shape proteomes

Peptides were reconstituted in 18μL buffer A* (0.1% FA, 2% ACN) fractionated on a 30 cm long 1.9 μm ReproSil C18 column (PepSep) using a 100min high-pH gradient. The concentration of Buffer B was increased from 3% to 30% in 45 min, to 40% in 12 min, to 60% in 5 min, to 95% in 10 min, kept constant for 10 min, reduced to 5% in 10 min and kept constant for 8 min. The eluted peptides were automatically collected into 48 fractions with a concatenation time of 90 seconds per fraction. The fractions were dried in a SpeedVac, reconstituted in 0.1% FA and directly loaded onto Evotips.

### Labelling of single-shape reference proteome

Peptides were reconstituted to 0.125 μg/μL in 60 mM triethylammonium bicarbonate buffer with 10% CAN, pH 8.5. The peptides were then dimethyl-labeled with 0.15% light formaldehyde (CH_2_O) and 0.023 M sodium cyanoborohydrate (NaBH_3_CN) for 1 hour at room temperature, quenched with 0.13% ammonia and acidified to 1% TFA. After drying in a SpeedVac, pellets were re-constituted in 100 μL buffer A, and desalted via 5 μg C18 columns on an AssayMap (Agilent) following the standard protocol. The resulting reference peptides were dried, and reconstituted to 1 ng/μL in buffer A.

### Peptide preparation of single shapes and dimethyl labeling for multiplexing

Peptides were prepared semi-automated on a Bravo pipetting robot (Agilent), similar as previously described ^11^. For this, plates were removed from the freezer and centrifuged. The wells were then washed on the robot with 28 μL of 100% acetonitrile and dried in a SpeedVac (Eppendorf) at 45 °C for 20 minutes. Shapes were then re-suspended in 6 μL of 80 mM triethylammonium bicarbonate buffer (pH 8.5, Sigma) with 0.013% dodecyl-β-D-maltoside (DDM, Sigma), and cooked for 30 minutes at 95 °C in a PCR cycler at a lid temperature of 110 °C. After addition of 1 μL of 80% acetonitrile (final concentration 10%), samples were incubated for another 30 minutes at 75 °C, cooled briefly, and 1 μL with 4 ng LysC and 6 ng trypsin was added. We digested the samples for 18 hours, and added 1 μL of either intermediate (CD_2_O) or heavy formaldehyde (^13^CD_2_O) to a final concentration of 0.15%. Without delay, either light (NaBH_3_CN) or heavy (NaBD_3_CN) sodium cyanoborohydrate were added to 0.023 M to retrieve Δ4 and Δ8 dimethyl labeled single-shape samples. The sealed plate was then incubated at room temperature for 1 hour, and the reaction was quenched to 0.13% ammonia, and acidified to 1% TFA.

### Peptide loading onto C-18 tips

C-18 tips (Evotip Pure, EvoSep, Denmark) were activated for 5 minutes in 1-propoanl, washed twice with 50 μL of buffer B (99.9% ACN, 0.1% FA), activated for 5 minutes in 1-propanol, and washed twice with 50 μL buffer A (0.1% formic acid). Single-shape samples were then loaded automatically with the Agilent Bravo robot into 30 μL buffer in the tip that was spun through the C-18 disk for a few seconds only. For loading, 10 μL of 1 ng/μL reference peptides (Δ0) were pipetted first, followed by Δ4, and Δ8 samples with the same tip. Wells were rinsed with 15 μL buffer A that were also loaded onto the tip. After peptide binding, the disk was further washed once with 50 μL buffer A, and then overlayed with 150 μL buffer A. All centrifugation steps were performed at 700xg for 1 minute, expect sample loading for 2 minutes.

For five shape proteomes, plates were treated as above, with the exception of lysis in 4.5 μL 60 mM triethylammonium bicarbonate buffer, pH 8.5 without DDM, and consecutive addition of 1 μL Lys-C and 1.5 μL trypsin to achieve the same digestion volume as above. Five-shape samples were not dimethyl labeled and multiplexed, but acidified directly after digest, and loaded manually onto Evotips following the protocol described above.

### LC-MS/MS analysis of five-shapes

Samples were measured with the Evosep One LC system (EvoSep) coupled to a timsTOF SCP mass spectrometer (Bruker Daltonics, US). The 30SPD (samples per day) method was used with the Evosep Performance column 15 cm, 150 μm ID (EV1137 EvoSep, Denmark) at 40°C inside a nanoelectrospray ion source (Bruker Daltonics, US) with a 10 μm emitter (ZDV Sprayer 10, Bruker Daltonics, US). The mobile phases were 0.1% FA in LC–MS-grade water (buffer A) and 99.9% ACN/0.1% FA (buffer B). We used a dia-PASEF method with 16 dia-PASEF scans separated into 4 ion mobility windows per scan covering an m/z range from 400 to 1200 by 25 Th windows and an ion mobility range from 0.6 to 1.6 Vs cm^-2^ (‘standard scheme’ ^14^). The mass spectrometer was operated in high sensitivity mode, with an accumulation and ramp time at 100 ms, capillary voltage set to 1750V and the collision energy as a linear ramp from 20 eV at 1/K_0_ = 0.6 Vs cm^-2^ to 59 eV at 1/K_0_ = 1.6 Vs cm^-2^.

### LC-MS/MS analysis of single shapes

Samples were measured with the Evosep One LC system (EvoSep) coupled to a timsTOF SCP mass spectrometer (Bruker Daltonics, US). The Whisper40 SPD (samples per day) method was used with the Aurora Elite CSI third generation 15 cm and 75 μm ID (AUR3-15075C18-CS IonOpticks, Australia) at 50 °C inside a nanoelectrospray ion source (Bruker Daltonics, US). The mobile phases were 0.1% formic acid in LC–MS-grade water (buffer A) and 99.9% ACN/0.1% FA (buffer B). The timsTOF SCP was operated with an optimal dia-PASEF method generated with our Python tool py_diAID ^24^. The method contained 8 dia-PASEF scans with variable width and 2 ion mobility windows per dia-PASEF scan, covering an m/z from 300 to 1200 and an ion mobility range from 0.7 to 1.3 Vs cm^-2^, as previously used on the same gradient and similar input material amount ^11^. The mass spectrometer was operated in high sensitivity mode, with an accumulation and ramp time at 100 ms, capillary voltage set to 1400 V and the collision energy as a linear ramp from 20 eV at 1/K_0_ = 0.6 Vs cm^-2^ to 59 eV at 1/K_0_ = 1.6 Vs cm^-2^.

The labeling efficiency was accessed on the same LC-MS/MS in dda-PASEF mode with ten PASEF scans per topN acquisition cycle. Singly charged precursors were excluded by their position in the m/z-ion mobility plane using a polygon shape, and precursor signals over an intensity threshold of 1,000 arbitrary units were picked for fragmentation. Precursors were isolated with a 2 Th window below m/z 700 and 3 Th above, as well as actively excluded for 0.4 minutes when reaching a target intensity of 20,000 arbitrary units. All spectra were acquired within a m/z range of 100 to 1700. All other settings were kept as described before.

### Spectral library generation

The spectral library was generated on five dda-PASEF single shots from 50 ng mouse reference peptide, using the same chromatography method as above. Spectra were search with FragPipe v18.0 ^25^ using MSFragger v3.5, Philosopher v4.4.0, and EasyPQP v0.1.32 against a mouse FASTA reference file with 55319 entries used throughout this study, excluding 50% decoys. Standard settings of the DIA_SpecLib_Quant workflow were used with the following exceptions: N-terminal and lysine mass shift of 28.0313 Da were set as fixed modifications, and methionine oxidation as variable modification. Carbamidomethylation was unselected as samples were not reduced and alkylated. One missed cleavage was accepted. The precursor charge ranged from 2 to 4. The peptide mass range was set to 300 to 1,800, and peptide length from 7 to 30. For DIA-NN compatibility, the column ‘FragmentLossType’ was removed in the output library file.

### Spectral search

All 263 files were search together in DIA-NN (version 1.8.1) ^15^ against the above-generated library, using a mass and MS1 mass accuracy of 15.0, scan windows of 9, and activated isotopologues, MBR, heuristic protein inference and no shared spectra, in single-pass mode. Proteins were inferred from genes. Library generation was set as ‘IDs, RT & IM profiling’, and ‘Robust LC (high precision)’ as quantification strategy. Dimethyl labeling at N-termini and lysins was set as fixed modification at 28.0313Da, and Δ4 or Δ8 were spaced 4.0251 Da or 8.0444 Da from the reference Δ0 ({--fixed-mod Dimethyl, 28.0313, nK} and {--channels Dimethyl, 0, nK, 0:0; Dimethyl, 4, nK, 4.0251:4.0251; Dimethyl, 8, nK, 8.0444:8.0444}). Additional settings were -- original-mods --peak-translation --ms1-isotope-quant --report-lib-info.

### Data analysis

To determine the quantities of the precursors in the DIA-NN report.tsv file, we utilized the Python-based RefQuant algorithm ^11^. In brief, RefQuant determines the ratio between target- and reference channel for each individual fragment ion and MS1 peak that is available. This gives a collection of ratios from which RefQuant estimates a likely overall ratio between target and reference. The ratio between target and reference was rescaled by the median reference intensity over all runs for the given precursor, thereby giving a meaningful intensity value for the target channel. The RefQuant quantification matrix was filtered for ‘Lib.PG.Q.Value’ < 0.01, ‘Q.value’ < 0.01 and ‘Channel.Q.Value’ < 0.15 and was then collapsed to protein groups using the MaxLFQ algorithm ^26^ as implemented in the R package iq (v. 1.9.6) ^27^ with median normalization turned off. Protein group data was then further analyzed in R v4.2.1 operating in RStudio v2022.07.2. Samples were excluded if the number of detected proteins was below 1.5 or above 3 standard deviations from the sample identification median, or within [806, 3362] identified proteins, resulting into a dropout of 8.9% (41 of 459 samples). Four additional samples were removed due to their outlier position on the PCA. After sample filtering, data was median normalized to a set of proteins that was quantified across all samples (100% completeness, 175 proteins), thus correcting for the dependency of protein numbers on shape size into account. For hepatocyte specific analysis, the arteriole proteomes were removed prior to normalization. We manually chose eight proteome classes for all comparative spatial analyses, and five classes for machine learning as a compromise between meaningful separation and having enough samples per class. Proteome bins were based on an equidistant split of PC1 Eigenvalues, distance classes accordingly on a split of portal over central vein distance ratios, and applied as indicated. Principle Component Analyses (PCA) were performed with the PCAtools v2.8.0 package. Limma v3.52.4 was used for statistical testing across proteome bins on a 50%-complete protein data matrix. ‘FDR’ was applied for multiple testing correction, and an FDR cutoff of 5% was considered significant. Heatmapping was performed with pheatmap 1.0.12, the completeness of the data matrix is indicated in the figure legends. Proteomic Gene Set Enrichment Analyses (GSEA) were done with WebGestalt 2019 ^28^ in an R environment using KEGG metabolic pathways as a library and an FDR threshold for reporting of 1. Significance was defined as FDR < 5%, and normalized enrichment scores are reported here. Urea cycle and peroxisomal fatty acid degradation proteins were manually curated. Normality was assessed with Shapiro-Wilk’s test, and *p* values were corrected for multiple testing and expressed as false discovery rate. Spatial data from xml files was plotted with the package sf v1.0-9. For comparisons to scRNAseq data, the dataset of Halpern *et al*. ^6^ was used, for which we binned the proteome data into nine bins guided by PC1 as described above. The GSEA was calculated with WebGestalt at a reporting and significance FDR cutoff of 5%, and KEGG, Reactome, and Gene Ontology Biological Processes as libraries.

### Image processing

Image data analysis was done in Python (3.8.11). Image shapes were extracted from the stitched tiles using Pillow (9.0.0). For each shape, the bounding box was calculated by taking the floor and ceiling of each shape coordinate and taking the maximum and minimum in x and y. The bounding rectangle was used to crop out the respective region of interest (ROI) of the image. For image with offset extraction, the center of each bounding rectangle was calculated and rounded to the next integer. An offset of 1,000 was added to each direction to additionally capture the surrounding environment, and the bounding box was highlighted. For composite images, each image was exported per channel with matplotlib (3.5.1), reloaded, merged with NumPy (1.4.2), and saved again. ImageJ was used to manually measure the distance of a shape to its proximal portal and central vein.

### Machine learning

For each shape and in all four channels (CFP, Alexa488, Alexa568, Alexa647), the mean, median, minimum, and maximum intensity of each bounding box were calculated, as well as the shape area. This feature list was saved with pandas (1.22.3). Proteomics data was clustered with a k-means algorithm into five clusters. Next, we used a supervised learning approach to classify the proteomic clusters based on the feature list. The training was performed using the scikit-learn package (1.0.2). Data (n=408) was randomly split in train and test datasets (split = 0.2). For classification, we used a RandomForest-Classifier (n_estimators=200) and achieved a testing accuracy of 0.90. To export probabilities, we used the predict_proba functionality of RandomForest. Diagnostic plots were generated using the Yellowbrick package (1.5). The random state was set to 23 for train/test-split and RandomForestClassifier.

## Data availability

Proteomics and imaging data will be available upon publication.

## SUPPLEMENTARY FIGURES

**Supplementary Fig. 1:**
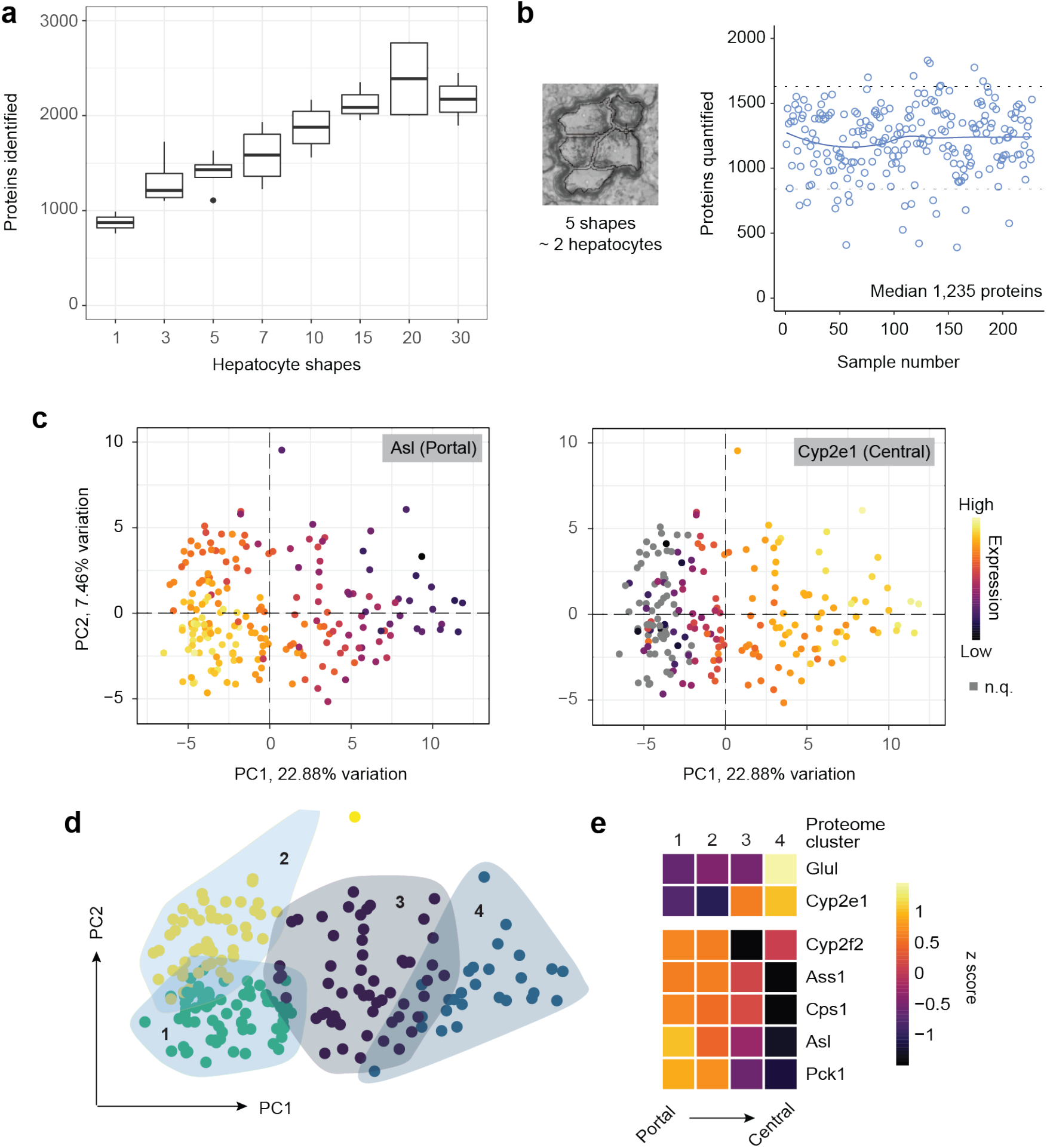
Five shape proteomes resolve liver zonation. **a**, Titration of number of shapes (10 μm thick) versus proteome depth achieved (n = 3). **b**, Protein numbers per five shapes across 230 samples. Line is a smoothing curve. **c**, Principal component analyses with a color overlay of two indicated zonation markers; n.q. not quantified. **d**, Unbiased *k* means clustering of all samples into four bins. **e**, Marker expression sorted by central (top) or portal (bottom) markers in the indicated *k* means clusters in d, expressed as z-score of log2 transformed protein abundances, and sorted according to summed zonal probability across all markers.

**Supplementary Fig. 2:**
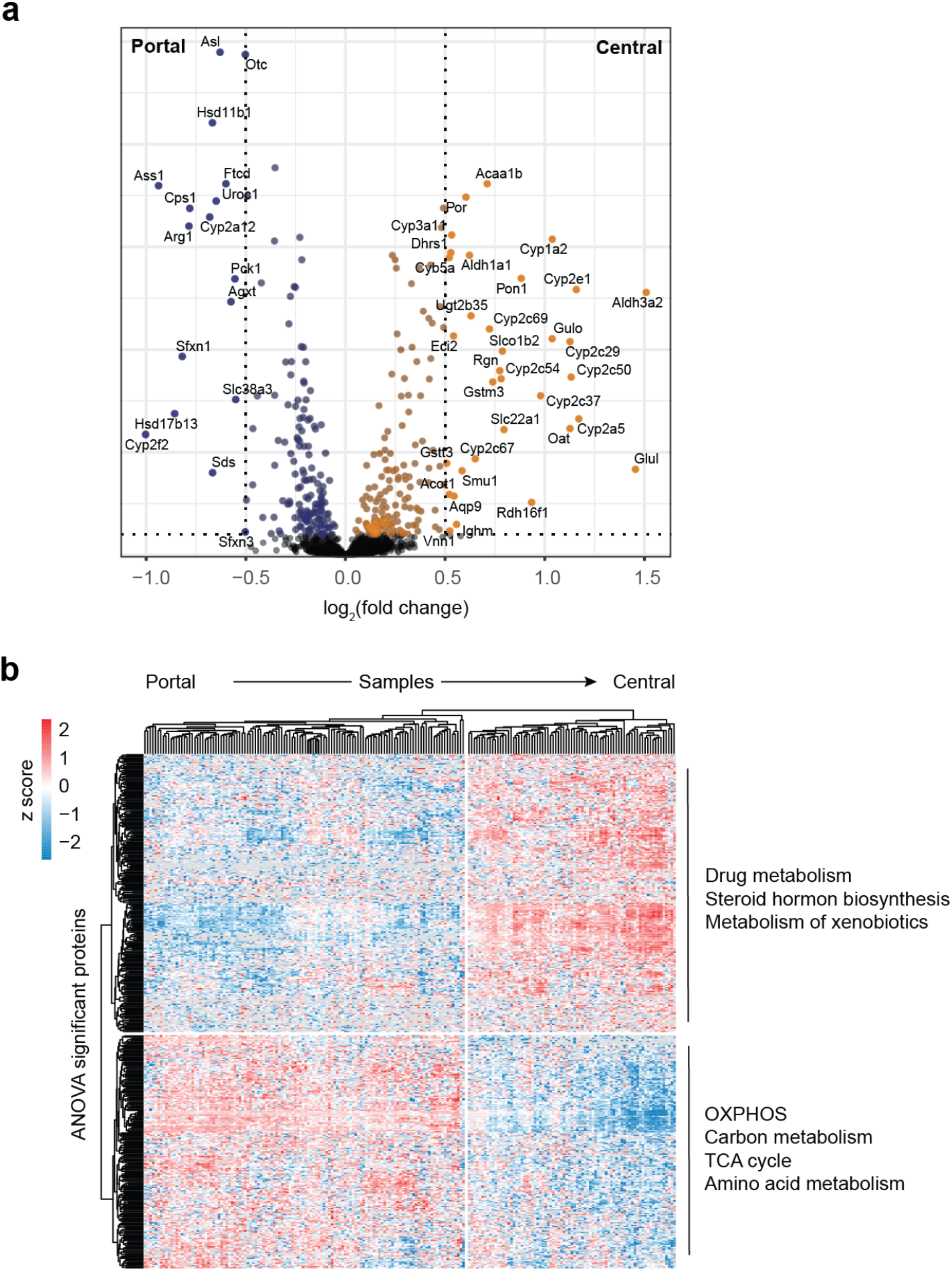
Statistical analysis of five shape proteomes. **a,** Volcano plot after an ANOVA over four sorted *k* means clusters (see Supplementary Fig. S1d). Statistically significant proteins (FDR < 0.05, n = 333 of 1652) with an absolute fold change of more than 0.5 are labeled. Colors indicate upregulation towards portal, or central zones. b, Overrepresentation analysis of statistically significant proteins in a. Samples are grouped into two bins depending on their visual expression profile on the heatmap. Euclidian clustering for both samples and proteins. Significant terms (FDR < 0.05) are presented on the right. Protein expression as z-score of log2 transformed protein abundances.

**Supplementary Fig. 3:**
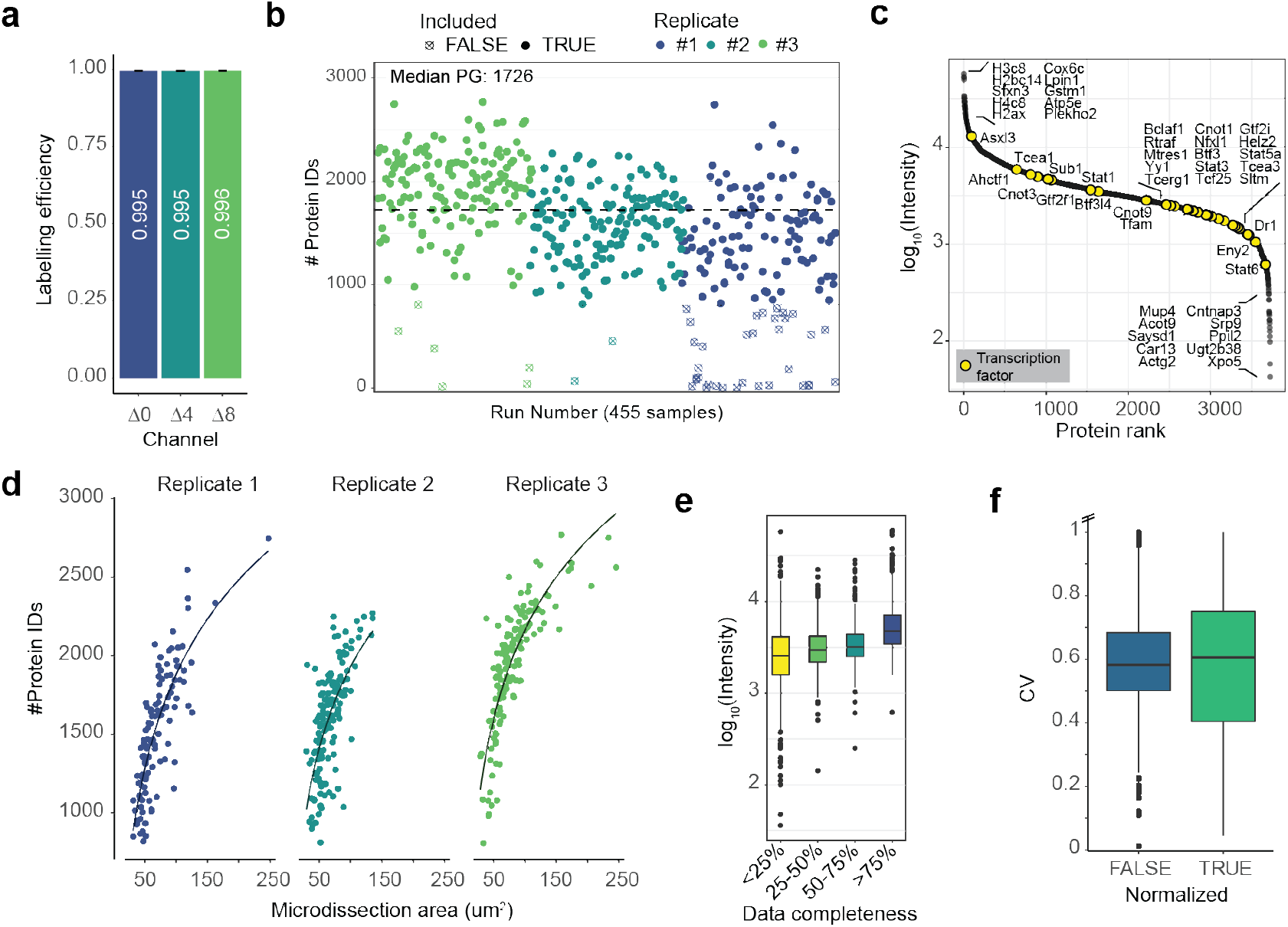
Performance overview of single shape proteomes. **a,** Labeling efficiency of 10 ng mouse liver peptide samples. Mean efficiency by intensity is stated in the bar (n = 5, mean +/-standard deviation). **b,** Number of proteins per sample (N = 455). The dotted line is the median, the fine pricked line is the sample exclusion cutoff of median minus 1.5 standard deviations. Samples were measured from left to right. Shape type indicates whether the samples was included for the final analysis. **c**, Association between the area of the cut shape, and number of proteins. Line is a log10 regression curve. **d,** Percentage of proteins quantified, binned into four groups, versus log10 transformed median intensities in the respective bin. **e,** Coefficient of variation (CV) prior and after complete proteome normalization (see Methods section). **f,** Proteins ranked by median intensity across all samples, versus median log10 transformed intensity. Top and bottom 10 proteins are indicated.

**Supplementary Fig. 4:**
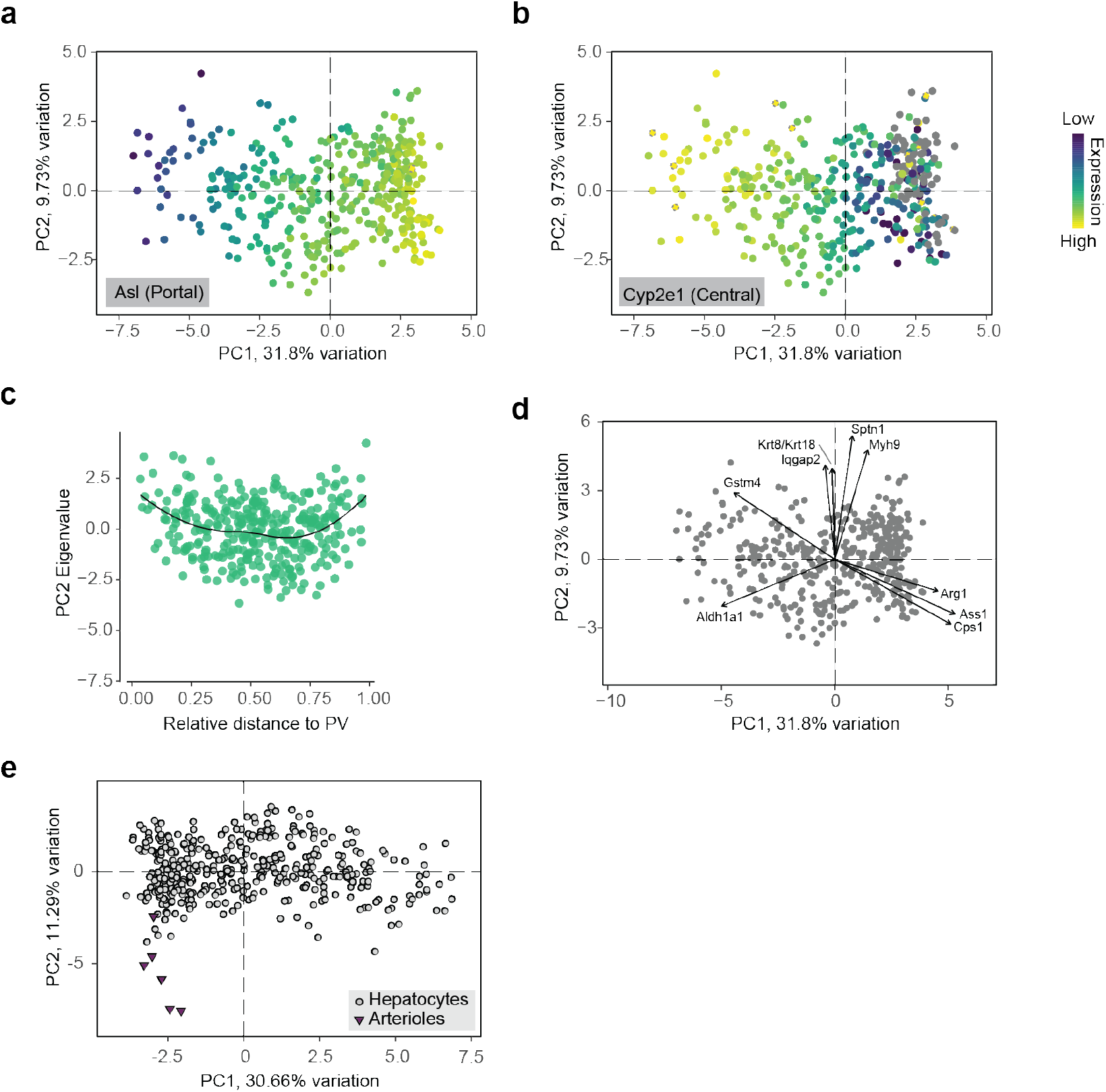
Dimensionality reduction of single shape data. **a,** Color overlay is expression level of the portal marker Asl, or **b,** the central marker Cyp2e1. **c,** Eigenvalues of PC2 versus measured distance ratio portal over central vein for all shapes. D, Top 10-leading edges as Eigenvectors (arrows) with proteins. E, Arterioles were cut as quality controls (see Methods section), and separate from hepatocytes on PC2 (n = 6).

**Supplementary Fig. 5:**
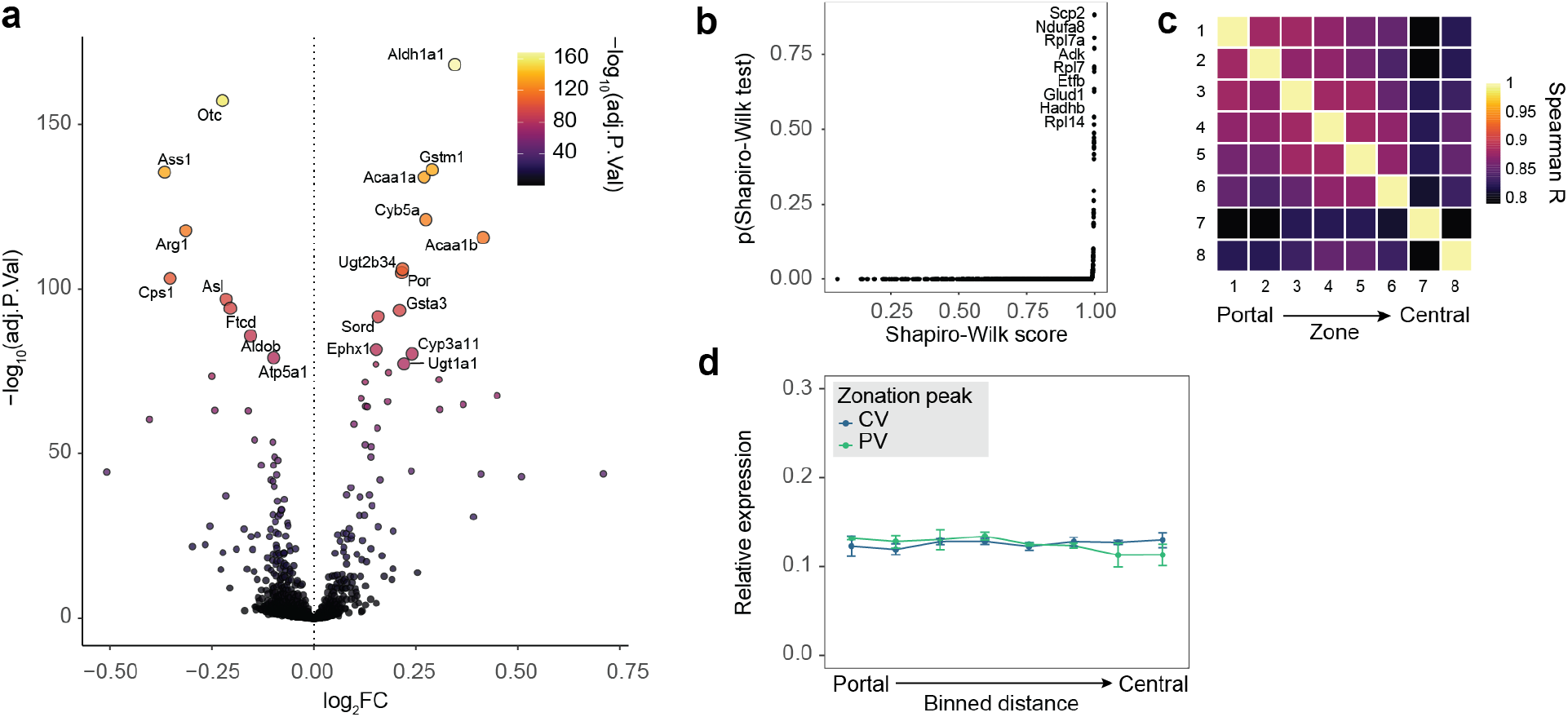
Functional analysis of single shape data. **a,** Volcano plot after ANOVA across eight PC1-guided proteome bins. Color overlay specifies adjusted *p* value, the top 20 significant proteins are labeled. b, Score and multiple testing adjusted *p* value of a Shapiro-Wilk normality test. Lowest proteins are labeled. c, Spearman correlation matrix with heatmap color overlay indicating Spearman’s R, comparing the eight PC1-guided proteome zones. d, Relative expression normalized to 1 for each contributing protein (n = 10) of the least significant Shapiro-Wilk hits in b, from portal to central distance-guided bins.

**Supplementary Fig. 6:**
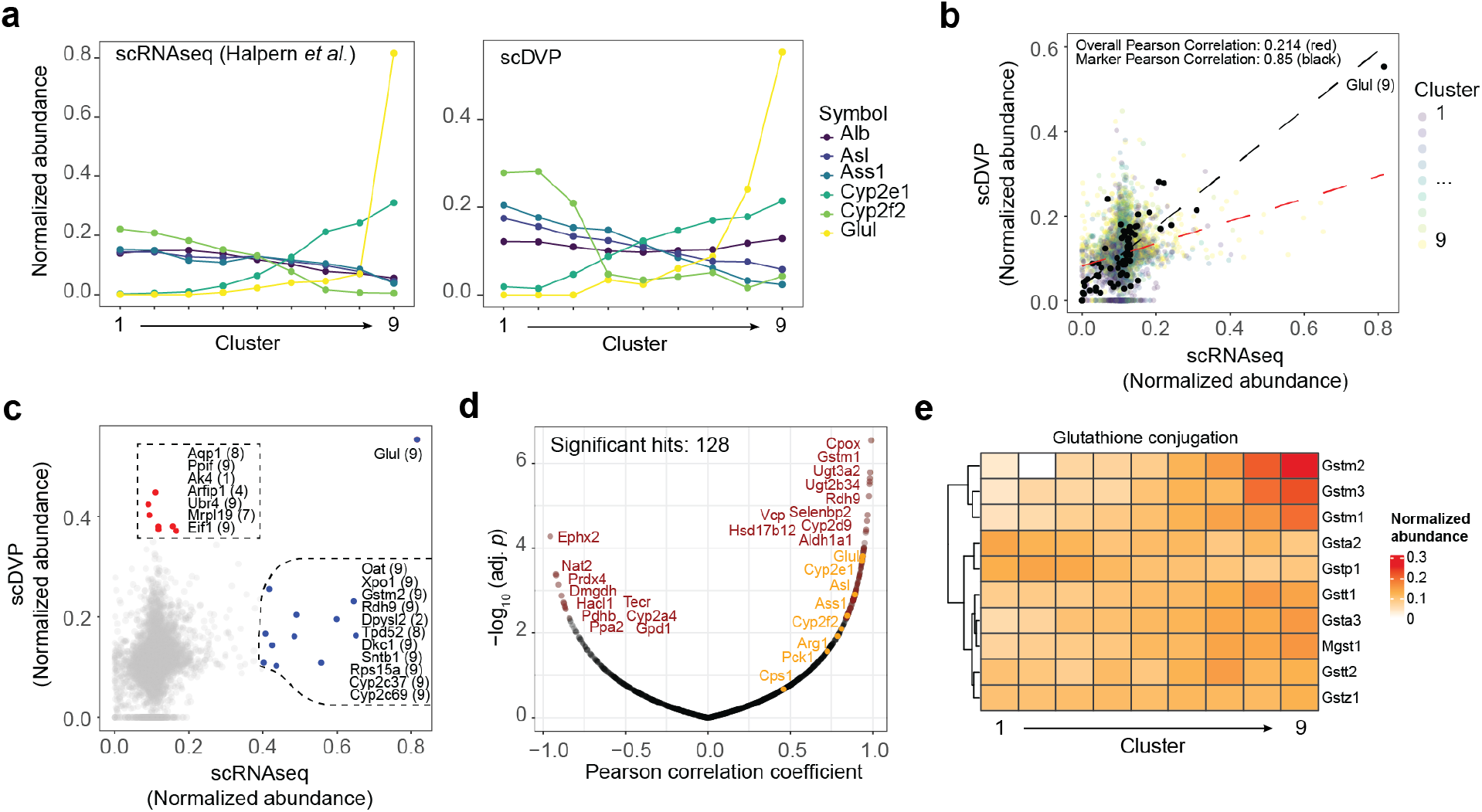
Comparison of scDVP and scRNAseq data. **a,** Abundance normalized to 1 across 9 bins in Halpern *et al.* ^6^ (left, marker expression guided bins), and this scDVP data (PC1-guided bins). **b,** Intensity correlation of all hits (opaque dots, color according to cluster) and markers (black dots). Linear regression as dashed line, with Pearson correlation coefficient given in the figure. One prominent hit marked with zone in brackets. **c,** As b, with outliers marked. Blue: Regulated on transcript level, red: regulated on protein level. Corresponding zone in brackets. **d,** Correlation coefficient for targets across all bins, with multiple testing adjusted *p* value. Top ten hits on either side are labeled in dark red, and marker proteins in orange. **e,** A significant hit after gene set enrichment analysis on Pearson correlation coefficient, with normalized abundance of protein levels as heatmap colors.

**Supplementary Fig. 7:**
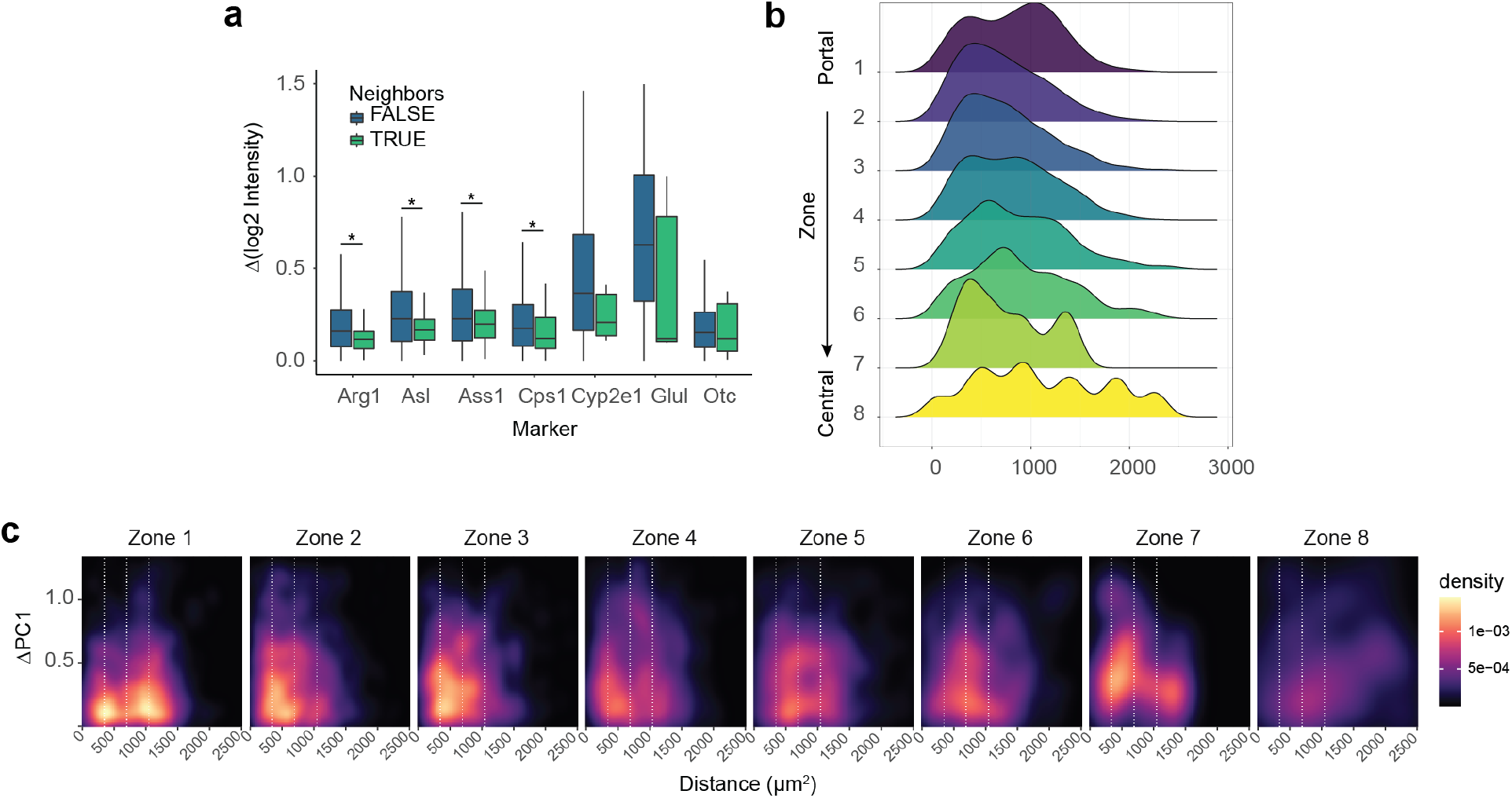
Neighborhood analysis of scDVP data. **a,** Intensity difference of directly adjacent neighbors (‘TRUE’) versus samples in the same proteome bin, but different lobules (‘FALSE’), for selected marker proteins. **b,** Measured distance of samples that belong to the same PC1-guided proteome bin. **c**, As in b, measured distance against difference in PC1 Eigenvalue, the color scheme represents data density. White dotted intersect at multiples of 350 μm.

**Supplementary Fig. 8:**
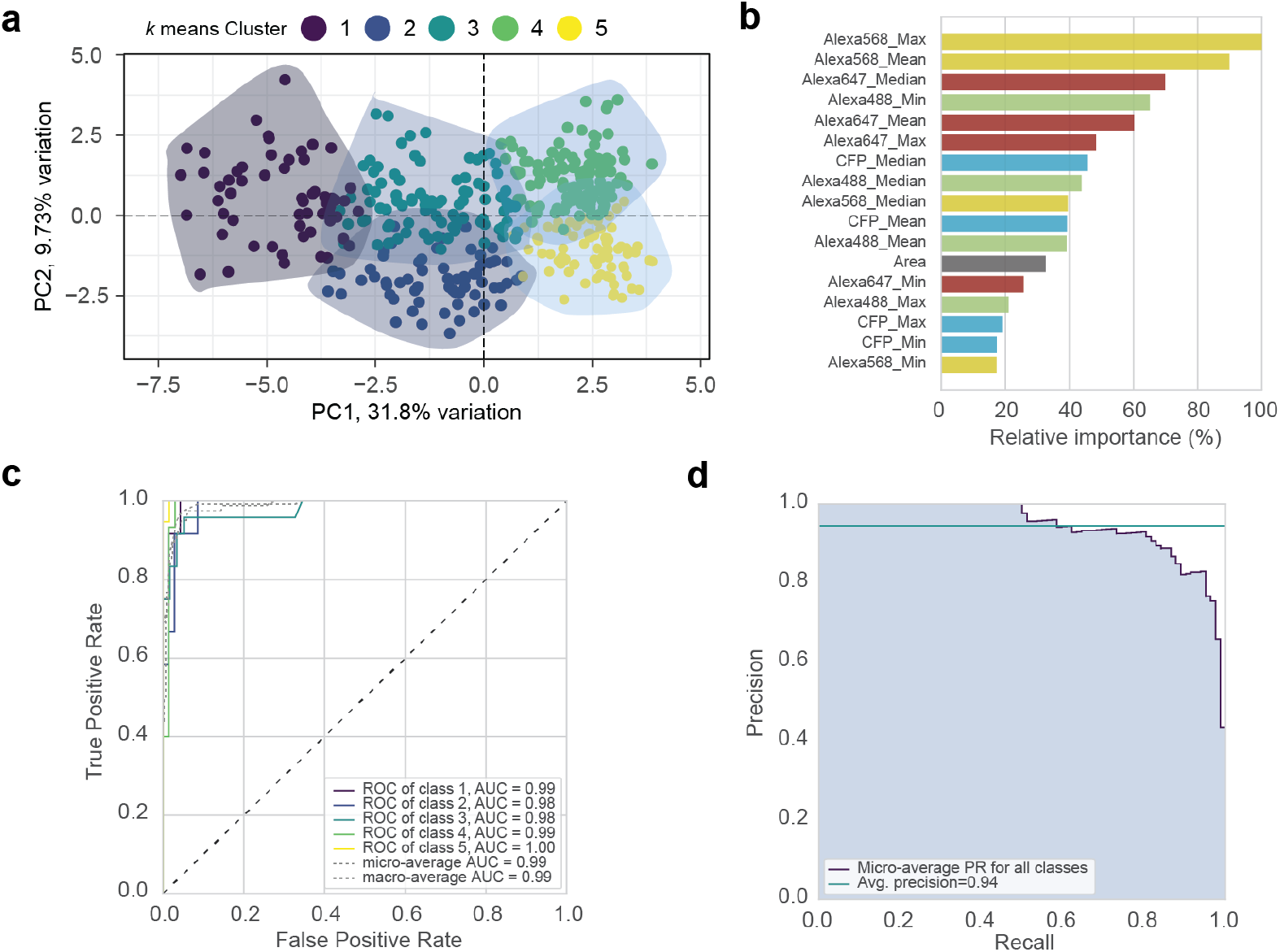
Machine learning (ML) accurately predicts proteome class. **a,** *k* means clustering, dividing all samples into five classes that inform the ML. **b**, Feature importance of the ML model, relative to the highest contributor. **c**, Receiver-Operating-Characteristics for each class. The individual Area Under the Curve (AUC) is given in the graph. **d**, Precision-recall-curve for the five classes.

