## Supplementary Figures S1 - S8 for "Spatial single-cell mass spectrometry defines zonation of the hepatocyte proteome"

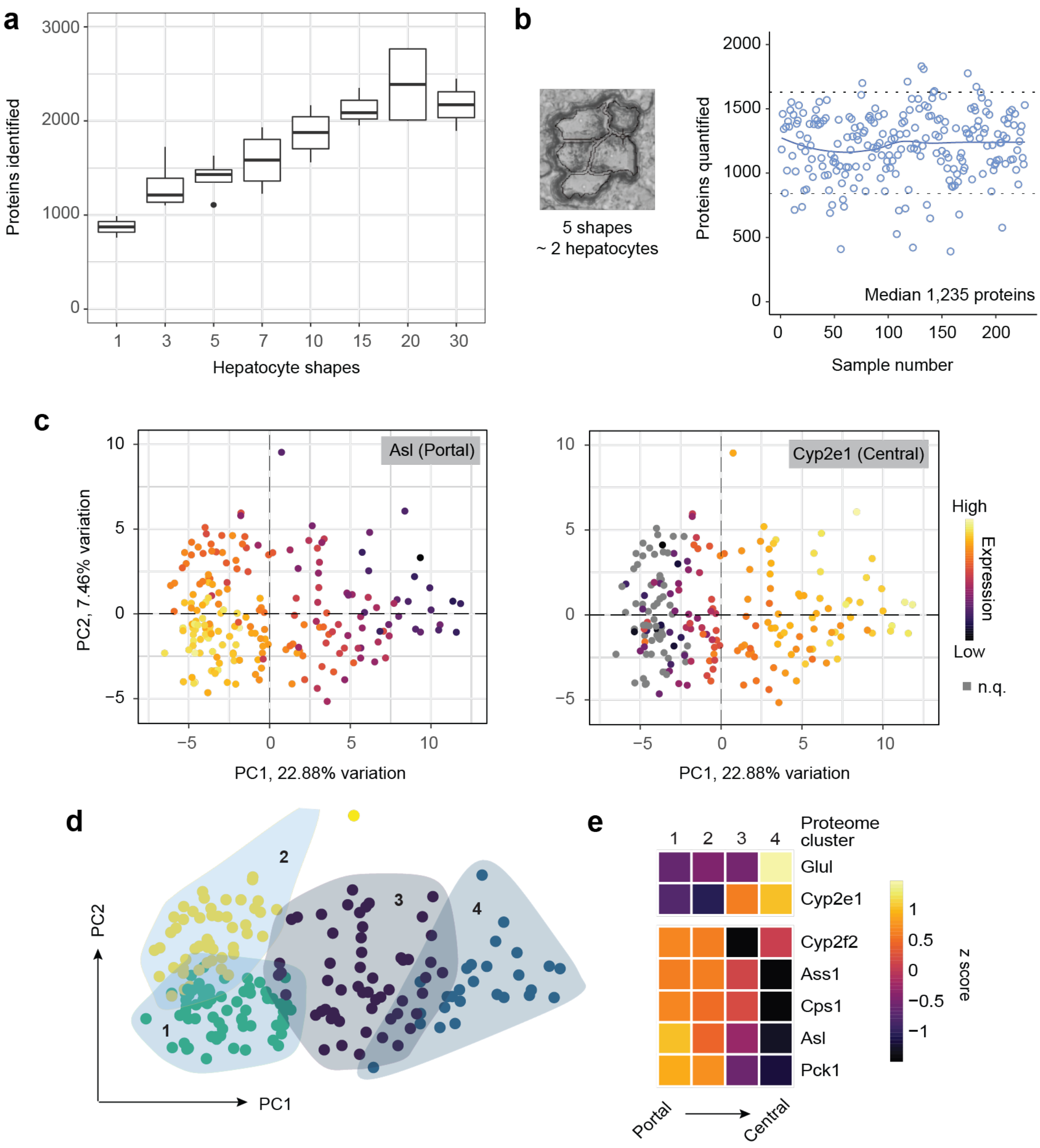


**scDVP, Supplementary Figure 1**

**Supplementary Fig. 1: Five shape proteomes resolve liver zonation. a**, Titration of number of shapes (10 µm thick) versus proteome depth achieved (n = 3). **b**, Protein numbers per five shapes across 230 samples. Line is a smoothing curve. **c**, Principal component analyses with a color overlay of two indicated zonation markers; n.q. not quantified. **d**, Unbiased *k* means clustering of all samples into four bins. **e**, Marker expression sorted by central (top) or portal (bottom) markers in the indicated *k* means clusters in d, expressed as z-score of log2 transformed protein abundances, and sorted according to summed zonal probability across all markers.

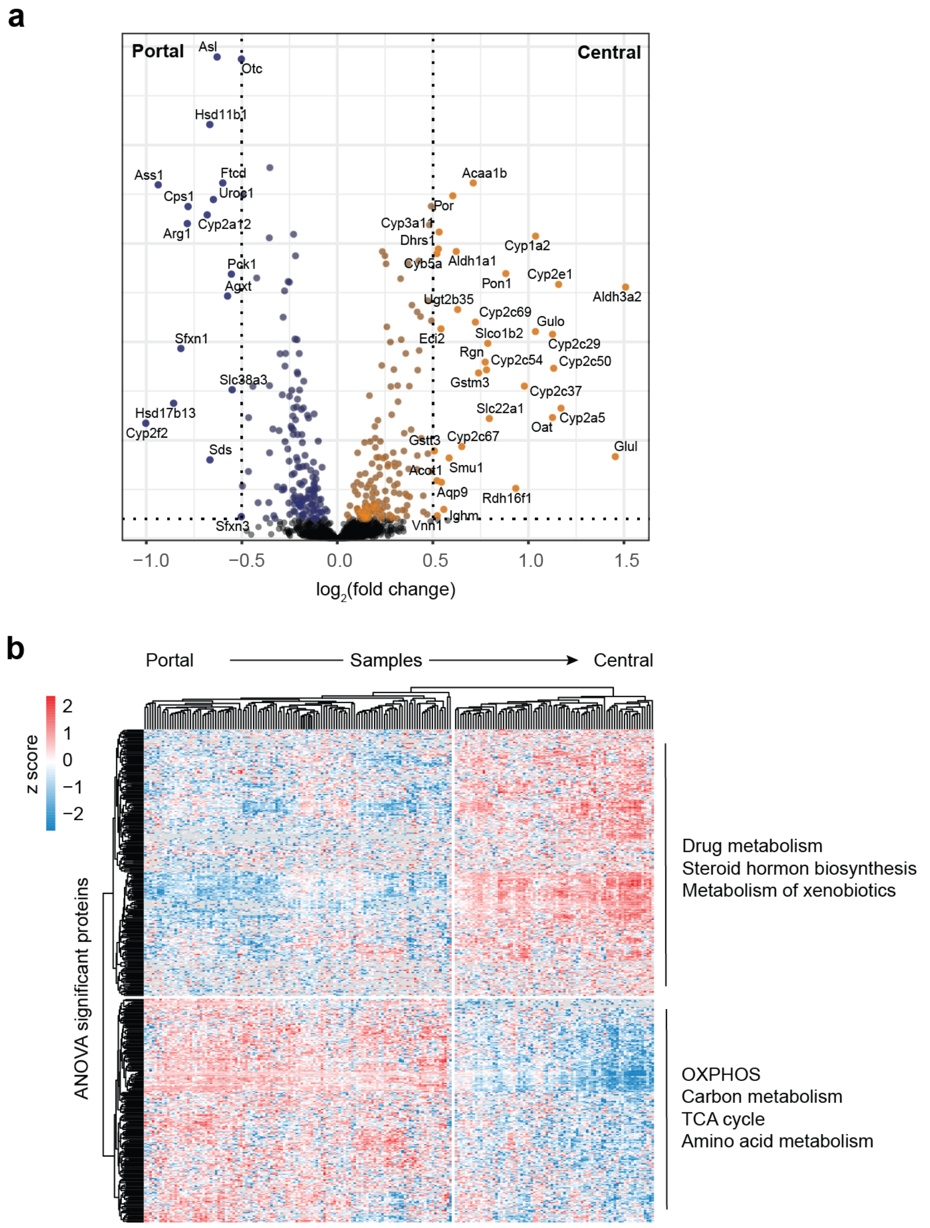


**scDVP, Supplementary Figure 2**

**Supplementary Fig. 2: Statistical analysis of five shape proteomes. a,** Volcano plot after an ANOVA over four sorted *k* means clusters (see Supplementary Fig. 1d). Statistically significant proteins (FDR < 0.05, n = 333 of 1652) with an absolute fold change of more than 0.5 are labeled. Colors indicate upregulation towards portal, or central zones. b, Overrepresentation analysis of statistically significant proteins in a. Samples are grouped into two bins depending on their visual expression profile on the heatmap. Euclidian clustering for both samples and proteins. Significant terms (FDR < 0.05) are presented on the right. Protein expression as z-score of log2 transformed protein abundances.

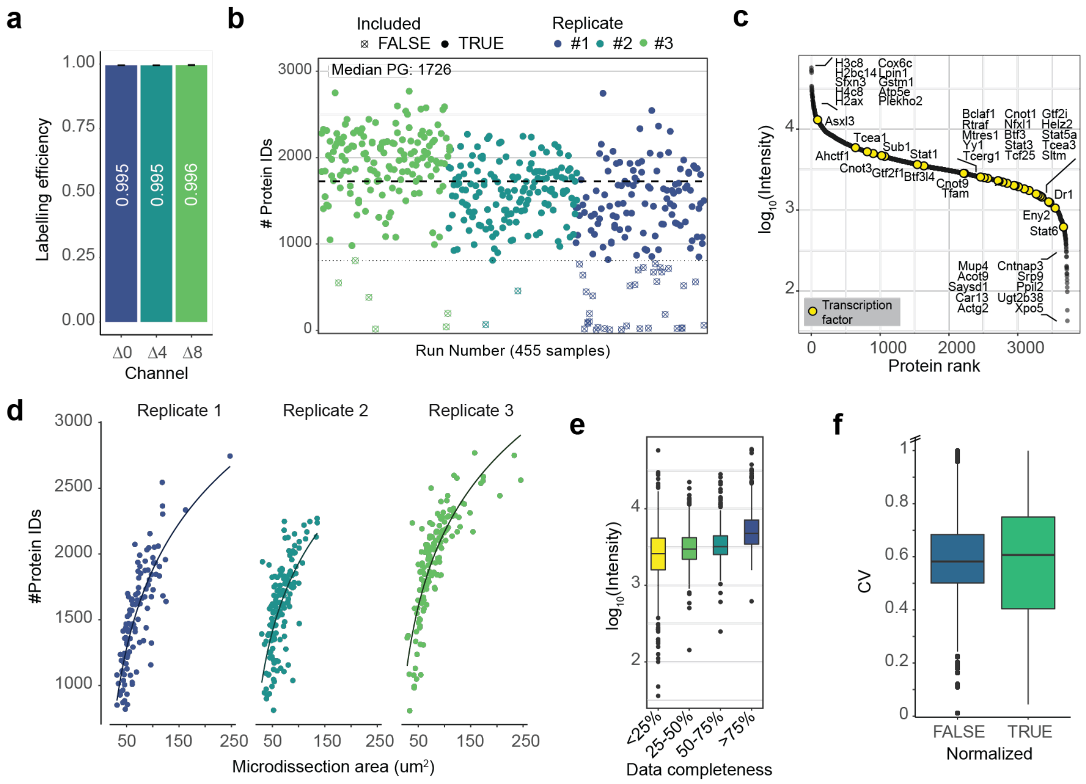

**scDVP, Supplementary Figure 3**

**Supplementary Fig. 3: Performance overview of single shape proteomes. a,** Labeling efficiency of 10 ng mouse liver peptide samples. Mean efficiency by intensity is stated in the bar (n = 5, mean +/- standard deviation). **b,** Number of proteins per sample (N = 455). The dotted line is the median, the fine pricked line is the sample exclusion cutoff of median minus 1.5 standard deviations. Samples were measured from left to right. Shape type indicates whether the samples was included for the final analysis. **c**, Association between the area of the cut shape, and number of proteins. Line is a log10 regression curve. **d,** Percentage of proteins quantified, binned into four groups, versus log10 transformed median intensities in the respective bin. **e,** Coefficient of variation (CV) prior and after complete proteome normalization (see Methods section). **f,** Proteins ranked by median intensity across all samples, versus median log10 transformed intensity. Top and bottom 10 proteins are indicated.

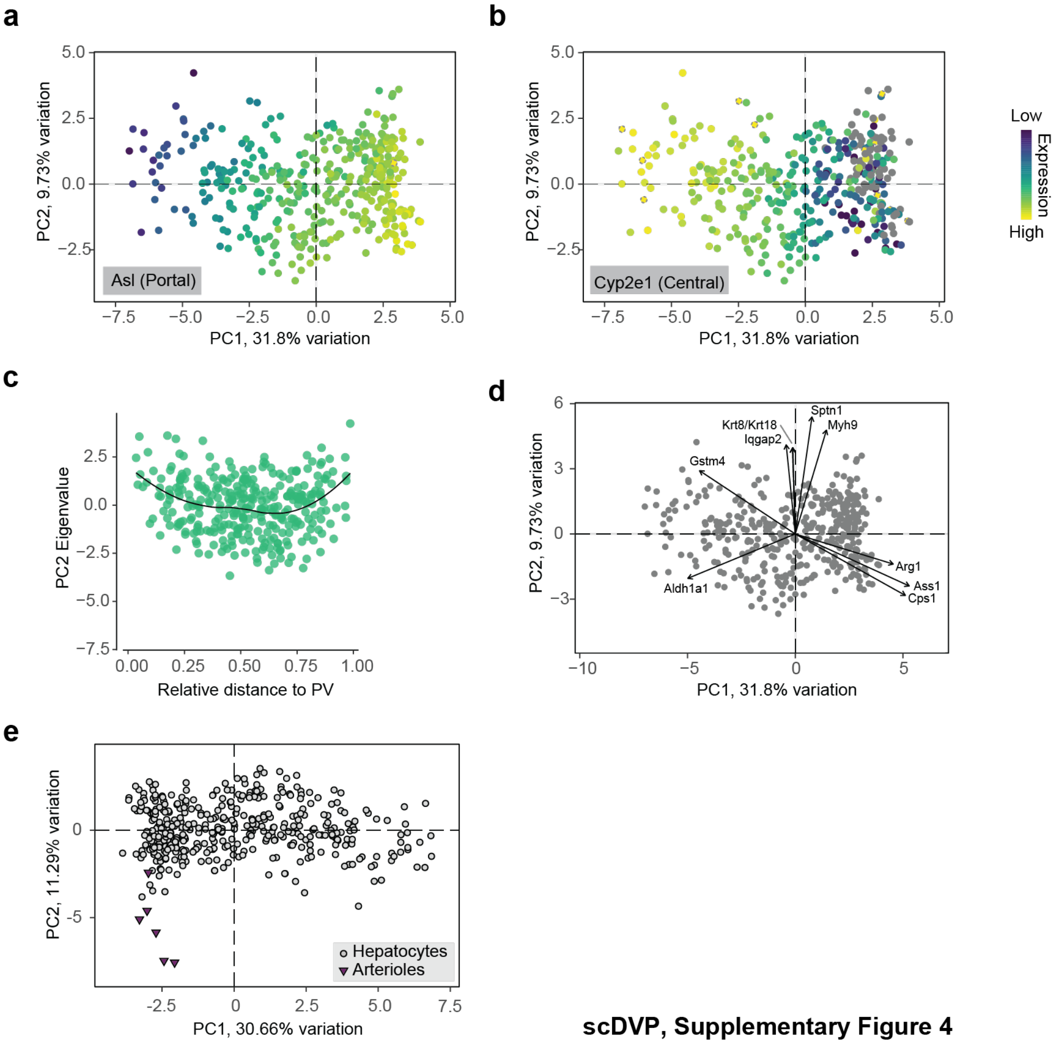


**scDVP, Supplementary Figure 4**

**Supplementary Fig. 4: Dimensionality reduction of single shape data. a,** Color overlay is expression level of the portal marker Asl, or **b,** the central marker Cyp2e1. **c,** Eigenvalues of PC2 versus measured distance ratio portal over central vein for all shapes. D, Top 10-leading edges as Eigenvectors (arrows) with proteins. E, Arterioles were cut as quality controls (see Methods section), and separate from hepatocytes on PC2 (n = 6).

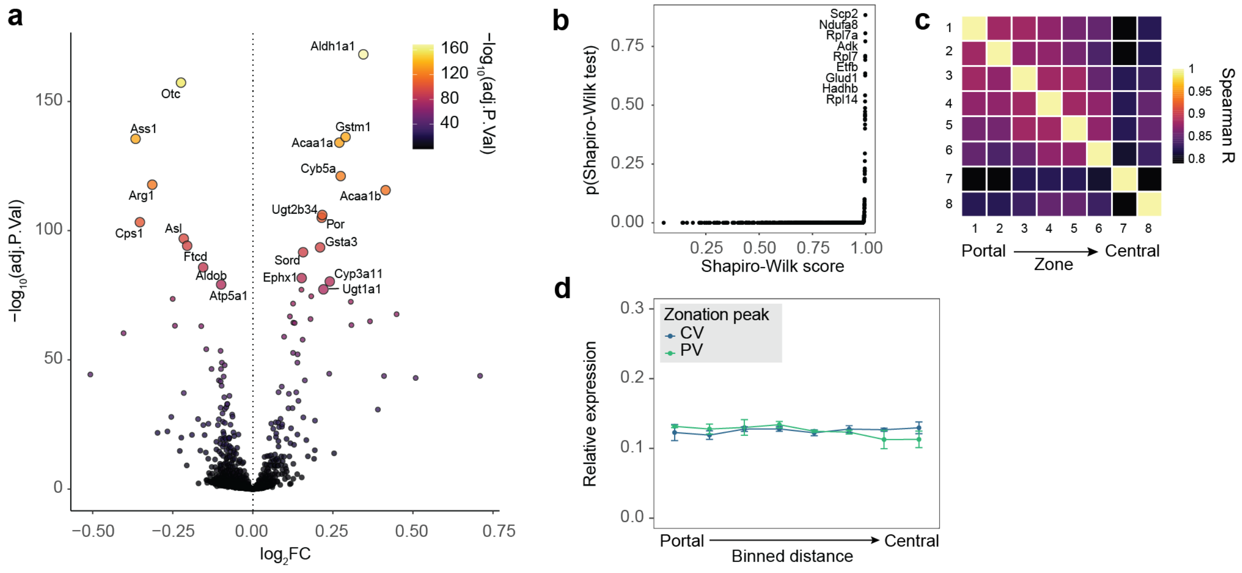


**scDVP, Supplementary Figure 5**

**Supplementary Fig. 5: Functional analysis of single shape data. a,** Volcano plot after ANOVA across eight PC1-guided proteome bins. Color overlay specifies adjusted *p* value, the top 20 significant proteins are labeled. b, Score and multiple testing adjusted *p* value of a Shapiro-Wilk normality test. Lowest proteins are labeled. c, Spearman correlation matrix with heatmap color overlay indicating Spearman's R, comparing the eight PC1-guided proteome zones. d, Relative expression normalized to 1 for each contributing protein (n = 10) of the least significant Shapiro-Wilk hits in b, from portal to central distance-guided bins.

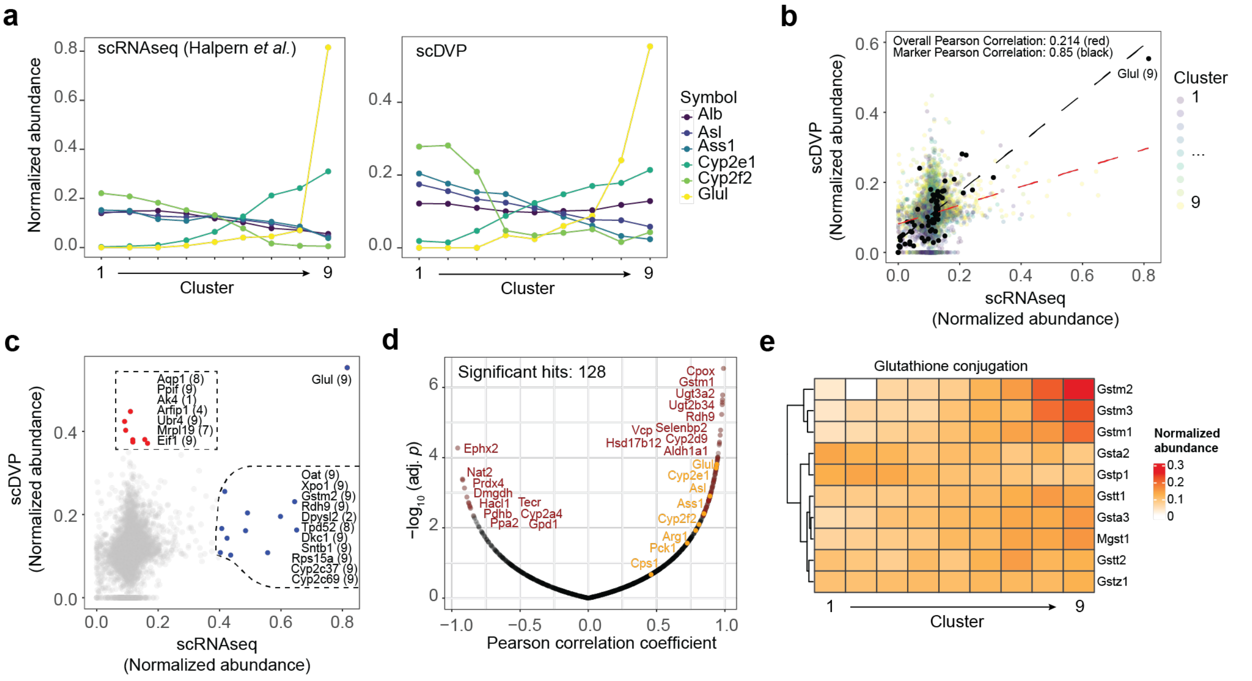


**scDVP, Supplementary Figure 6**

**Supplementary Fig. 6: Comparison of scDVP and scRNAseq data. a,** Abundance normalized to 1 across 9 bins in Halpern *et al.* [6] (left, marker expression guided bins), and this scDVP data (PC1-guided bins). **b,** Intensity correlation of all hits (opaque dots, color according to cluster) and markers (black dots). Linear regression as dashed line, with Pearson correlation coefficient given in the figure. One prominent hit marked with zone in brackets. **c,** As b, with outliers marked. Blue: Regulated on transcript level, red: regulated on protein level. Corresponding zone in brackets. **d,** Correlation coefficient for targets across all bins, with multiple testing adjusted *p* value. Top ten hits on either side are labeled in dark red, and marker proteins in orange. **e,** A significant hit after gene set enrichment analysis on Pearson correlation coefficient, with normalized abundance of protein levels as heatmap colors.

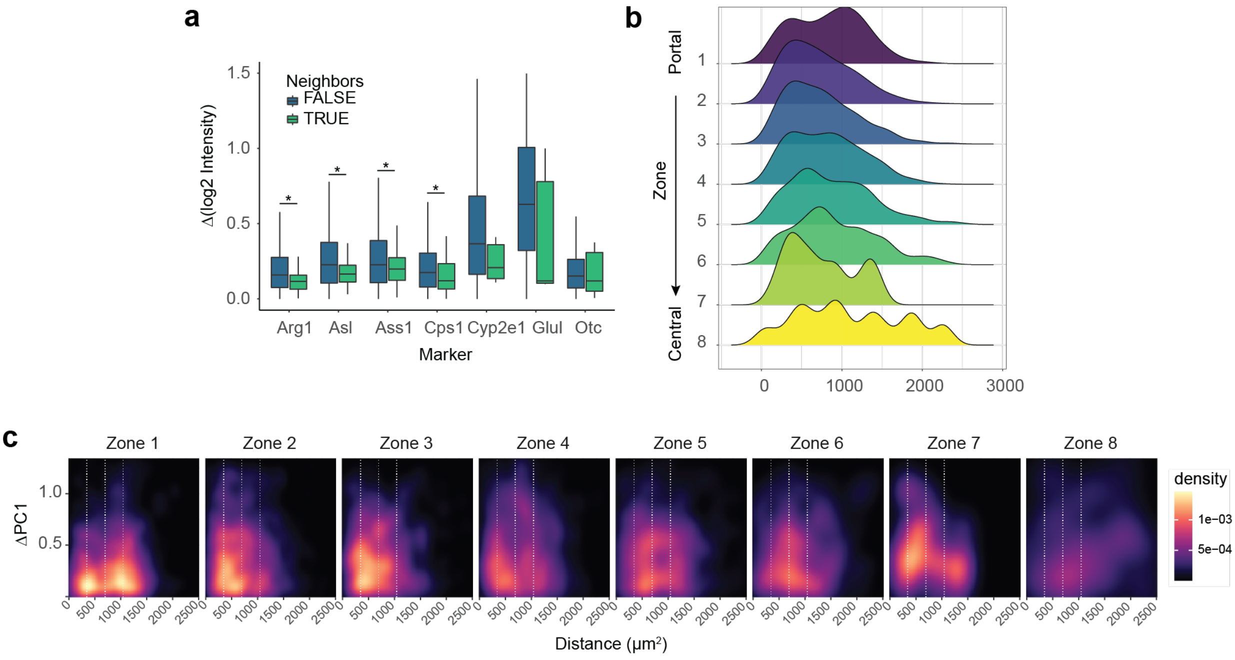


**scDVP, Supplementary Figure 7**

**Supplementary Fig. 7: Neighborhood analysis of scDVP data. a,** Intensity difference of directly adjacent neighbors ('TRUE') versus samples in the same proteome bin, but different lobules ('FALSE'), for selected marker proteins. **b,** Measured distance of samples that belong to the same PC1-guided proteome bin. **c**, As in b, measured distance against difference in PC1 Eigenvalue, the color scheme represents data density. White dotted intersect at multiples of 350 µm.

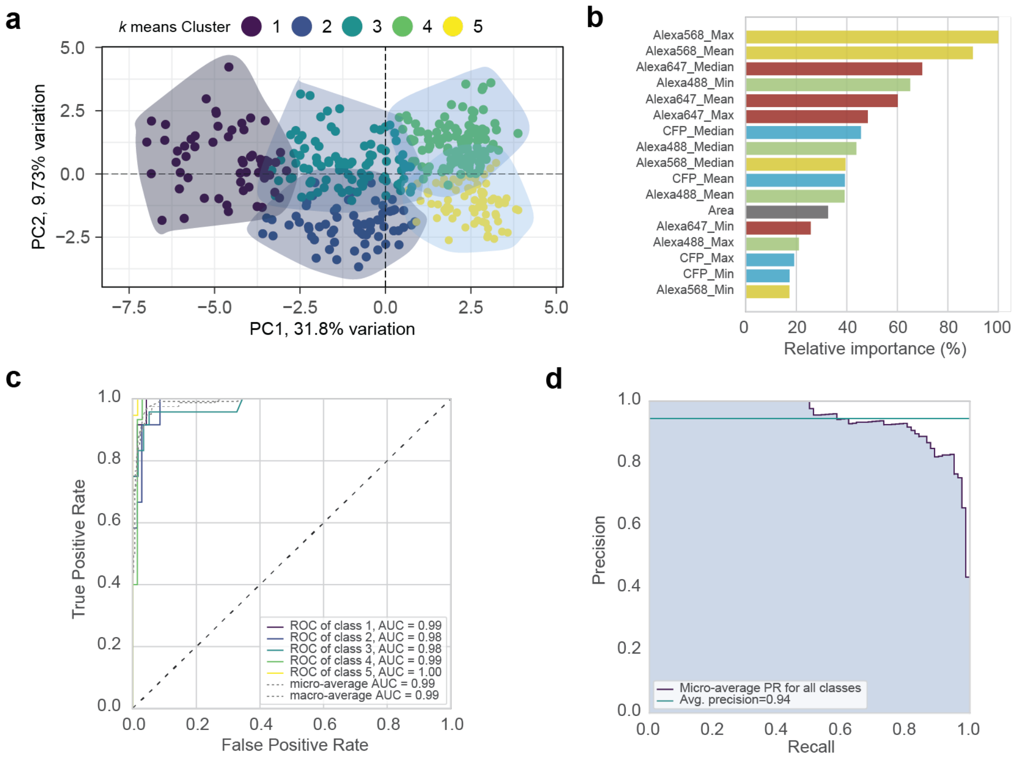


**scDVP, Supplementary Figure 8**

**Supplementary Fig. 8: Machine learning (ML) accurately predicts proteome class. a,** *k* means clustering, dividing all samples into five classes that inform the ML. **b**, Feature importance of the ML model, relative to the highest contributor. **c**, Receiver-Operating-Characteristics for each class. The individual Area Under the Curve (AUC) is given in the graph. **d**, Precision-recall-curve for the five classes.
